# *ClassifyMe*: a field-scouting software for the identification of wildlife in camera trap images

**DOI:** 10.1101/646737

**Authors:** G Falzon, C Lawson, K-W Cheung, K Vernes, GA Ballard, PJS Fleming, AS Glen, H Milne, AT Mather-Zardain, PD Meek

## Abstract

We present *ClassifyMe* a software tool for the automated identification of animal species from camera trap images. *ClassifyMe* is intended to be used by ecologists both in the field and in the office. Users can download a pre-trained model specific to their location of interest and then upload the images from a camera trap to a laptop or workstation. *ClassifyMe* will identify animals and other objects (e.g. vehicles) in images, provide a report file with the most likely species detections and automatically sort the images into sub-folders corresponding to these species categories. False Triggers (no visible object present) will also be filtered and sorted. Importantly, the *ClassifyMe* software operates on the user’s local machine (own laptop or workstation) not via internet connection. This allows users access to state-of-the-art camera trap computer vision software *in situ*, rather than only in the office. The software also incurs minimal cost on the end-user as there is no need for expensive data uploads to cloud services. Furthermore, processing the images locally on the users’ end-device allows them data control and resolves privacy issues surrounding transfer and third-party access to users’ datasets.

## Introduction

Passive Infrared sensor activated cameras, otherwise known as camera traps, have proved to be a tool of major interest and benefit to wildlife management practitioners and ecological researchers (Meek et al. 2014a;b). Camera traps are used for a diverse array of purposes including presence-absence studies (Khorozyan, Malkhasyan & Abramov 2008; Gormley et al. 2011; Ramsey, Caley & Robley 2015), population estimates (Karanth 1995; Trolle & Kéry 2003; Jackson et al. 2006; Gowen and Vernes 2014), animal behaviour studies (Vernes, Smith & Jarman 2014; Vernes & Jarman 2014; Vernes et al. 2015); and species interactions studies (Harmsen et al. 2009; Linkie & Ridout 2011; Meek, Zewe & Falzon 2012). A comprehensive discussion of applications of camera trap methodologies and applications are described in sources including O’Connell et al. (2011); Meek et al. (2014a); Rovero and Zimmerman (2016). The capacity of camera traps to collect large amounts of visual data provides unprecedented opportunity for remote wildlife observation, however these same datasets incur a large cost and burden as image processing can be time consuming (Meek et al. 2014b; Tack et al. 2016). The user is often required to inspect, identify and label tens-of-thousands of images per deployment; dependent on the number of camera traps deployed. Large scale spatio-temporal studies may involve 10-100s of cameras deployed consecutively over months to years and the image review requirements are formidable and resource intensive. Numerous software packages have been developed over the last 20 years to help with analysing camera trap image data (see Young, Rode-Margono & Amin 2018) but these methods often require some form of manual image processing. Automation in image processing has been recognised internationally as a requirement for progress in wildlife monitoring (Meek et al. 2014 a;b) and this has become increasingly urgent as camera trap deployment has grown over time.

Camera trap image interrogation can be tackled in different ways including; a) processing images by paid staff, b) use of the crowd-sourcing internet marketplace, c) recruiting volunteers such as citizen science programs or d) limiting the design of studies to ensure that data issues are manageable. Using staff requires access to sufficient budget and capable personnel and represents an expensive use of valuable resources in terms of both time and money. The quality of species identification is likely to be high but the time of qualified staff is otherwise lost for other tasks such as field work and data interpretation. Using the crowd-sourced internet market involves payment to commercial providers to out-source this task. A prominent example of such services is the *Amazon Mechanical Turk* (https://www.mturk.com/) whereby practitioners can submit their data to the service for processing for a fee. Tasks like species identification, also known as Human Intelligence Tasks (HIT’s), are widely used in commercial settings for the annotation of images. However, there are inherent biases between individuals with different expertise and aspirations often resulting in errors.

Crowd-sourcing involves the use of volunteers to provide image annotation services. Costs are comparatively low to that of employing staff but there a number of important limitations and considerations when designing machine learning algorithms in the presence of uncertainty in label veracity (Zhang, Wu & Sheng 2016). Species identification results could be poor as it might require specialised knowledge of the field. The suitability of this approach for sensitive ecological datasets (e.g. involving threatened or endangered species) is also an important consideration as is privacy laws and constraints (Meek and Butler 2014). There is also limited control and knowledge of where the data is transmitted and who accessed the data along with their locations and the storage of data records by service providers. Moreover, the potentially high cost of image data transfer over internet connections and issues around minimum wage and other ethical critiques of such services are controversial.

The use of volunteers or citizen scientists has proved effective in the field of camera trapping, notably via *TEAM Network* (Ahumada et al 2011) and the *Snapshot Serengeti* project (Swanson et al. 2015). In the context of camera trap image review, citizen science involves the review of camera trap imagery via a website. Images are randomly displayed along with species identification libraries and a visual identification guidebook. This approach is reportedly effective (Swanson et al. 2015) with iconic species but for some taxa human identification has been shown to be problematic (Meek, Vernes & Falzon 2013). Other challenges with the citizen science approach include access to the necessary software infrastructure (although services such as Zooinverse https://www.zooniverse.org now exist) along with privacy laws and related issues. Meek and Zimmerman (2016) discuss the challenges of using citizen science for camera trap research and managing such teams along with the data can incur enormous costs to the researchers. Furthermore, the researcher has limited control over the time taken to complete the review of the camera trap image datasets because it relies on the goodwill of the volunteers.

Limiting the design of studies to minimise analysis time by reducing the number of camera traps deployed, reviewing data for the presence of select species only or evaluating only a proportion of the available data and archiving the remainder is an unpalatable option. This approach does not fulfil the tests of scientific rigour (Claridge and Paull 2014; Swann & Perkins 2014) and constrains the value of such research findings for improving biodiversity management outcomes.

To overcome the limitations of approaches outlined above, including human error and operator fatigue, we have utilised computer science to develop automated labelling. As well as being able to validate results, key strengths of this approach, compared to existing options, include it being consistent, comparatively fast, standardised, and relatively free from biases associated with anthropomorphic values and operator fatigue. Advances in computer vision have been pronounced of recent years with successful demonstrations of image recognition in fields as diverse as autonomous cars, citrus tree detection from drone imagery and identification of skin cancer (Zhang et al. 2018; Csillik et al. 2018; Esteva et al. 2017). Recent work has also demonstrated the feasibility of Deep Learning approaches for species identification in camera trap images (Norouzzadeh et al. 2018) although it is worth noting that such algorithms have been used in prototype software for this purpose since at least 2015 in projects such as *Wild Dog Alert* (https://invasives.com.au/research/wild-dog-alert/) (Meek et al. in press) building on earlier semi-automated species recognition algorithms (Falzon, Meek & Vernes 2014). The practical benefit of this research for end-users has been limited because they cannot access software to automatically process camera trap images.

We therefore developed *ClassifyMe* as a software tool to reduce time and costs of image processing. The *ClassifyMe* software is designed to be used in the field on a laptop. This is a challenging requirement for a software application because it is required to operate across diverse computer hardware and software configurations while providing the end-user with a high-level of control and independence of their data. To elaborate on how we tackle these issues we outline the general structure and operation of *ClassifyMe* and provide an evaluation of its performance using an Australian species case study.

## Software Design

### Workflow

The software is developed so it can be installed on individual computers under an End User Licence Agreement (EULA). The intent is that the user will upload an SD card of camera trap images, select the relevant model and then run *ClassifyMe* on this dataset to automatically identify and sort the images (Fig 1).

**Figure 1.**
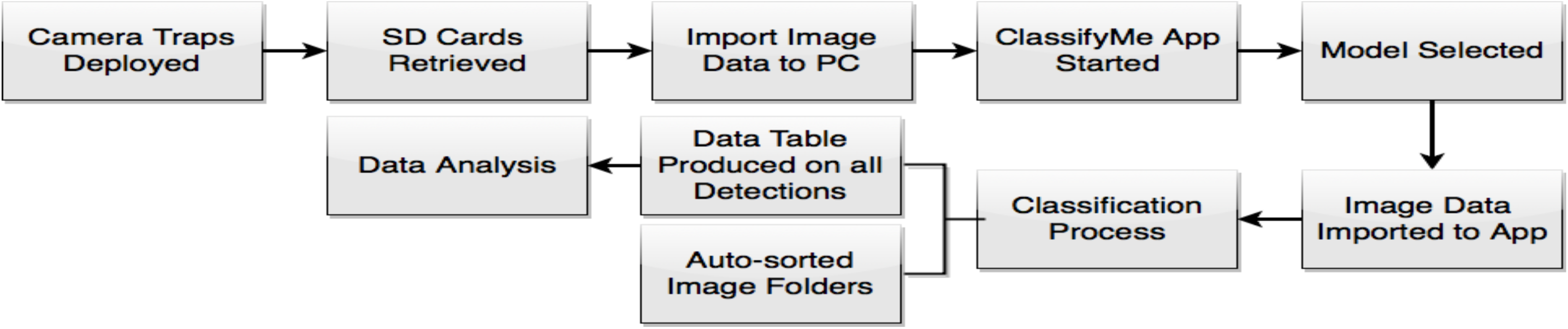
The data collection-analysis pipeline using the *ClassifyMe* software.

Processing camera trap images on a user’s own machine provides a high level of control on the use and access to the data. There are ongoing concerns around the sharing, privacy and security of using a cloud service or similar. Furthermore, *ClassifyMe* avoids the need for the user to upload their data to a cloud service which can be prohibitive in terms of accessibility, time and cost. *ClassifyMe* adopts a ‘tethered’ service approach whereby the user needs only intermittent internet access (every 3 months) to verify security credentials to ensure continued access to the software. The ‘tethered’ service approach was adopted as a security mechanism to obstruct misuse and unauthorised proliferation of the software for circumstances such as poaching. A practitioner can therefore validate security credentials and download the appropriate regional identification model (e.g. New England model) prior to travel into the field. When in the field, *ClassifyMe* can be used to evaluate deployment success (e.g. after several weeks of camera trap data collection) and can be used in countries with limited or no internet connectivity. Validation services are available for approved users (e.g. ecology researchers or managers) who require extensions of tethered renewal period.

### Software Design Attributes

The software design and stability of *ClassifyMe* was complicated by our choice to operate solely on the user’s computer. As such the software is capable of operating on a plethora of different operating systems and hardware designs. To limit stability issues *ClassifyMe* is designed to work with Windows 10™ as this operating system is widely used by field ecologists. Different hardware options are supported including CPU-only and GPU, the models used by *ClassifyMe* are best supported by NDVIDIA GPU hardware and as a result, users with this hardware will experience substantially faster processing times (up to 20 times faster per dataset).

The ‘tethered’ approach and corresponding application for software registration might be viewed as an inconvenience by some users. However, these components are essential security aspects of the software. The *ClassifyMe* software is a decentralised system; individual users access a web site, download the software and the model and then process their own data. The *ClassifyMe* web service doesn’t see the user’s end data and without the registration and ‘tethering’ process the software could be copied and redistributed in an unrestricted manner. When designing *ClassifyMe*, the authors were in favour of free, unrestricted software which could be widely redistributed. During the course of development, it occurred to the team the software was also at risk of misuse. In particular, *ClassifyMe* could be used to rapidly scan camera trap images whilst in field to detect the presence of particular species such as African elephants which are threatened by poaching (Bennett 2015). To address this concern a host of security features were incorporated into *ClassifyMe*. These features range from legal provisions on the software licence, through to a user validation and certification process and extensive undisclosed software security features. Disclosed security features include tethering and randomly generated licence keys and facilities to ensure that *ClassifyMe* is used only on the registered hardware and unauthorised copying is prevented. In the event of a breach attempt, a remote shutdown of the software is initiated.

All recognition models are restricted, and approval is issued to users on a case-by-case basis. This security approach is implemented in a privacy-preserving context. The majority of security measures involve hidden internal logic along with security provisions of the communications with the corresponding *ClassifyMe* web service https://classifymeapp.com/ (to ensure security of communications with the end user and their data). Information provided by the user and the corresponding hardware ‘fingerprinting’ identification is performed only with user consent and all information is stored on secured encrypted databases.

A potential disadvantage of the local processing approach adopted by *ClassifyMe* is that user’s software resources are utilised, which potentially limits the scale and rate of data processing. An institutional cloud service for instance can auto-scale (once the data is uploaded) to accommodate data sets from hundreds of camera trap SD card simultaneously. In contrast, the *ClassifyMe* user will only be able to only process one camera trap dataset at a time. The *ClassifyMe* user will also have to implement their own data record management system, there is no database system integrated within *ClassifyMe* which has the benefit of reducing software management complexity for end users but the disadvantage of not providing a management solution for large volumes of camera trap records. *ClassifyMe* is designed simply to review camera trap data for species identification, auto-sort of images and export of the classifications (indexed to image) to a csv file.

### Graphical User Interface

When *ClassifyMe* is initiated the main components consist of: a) an image banner which displays thumbnails of the camera trap image dataset, b) a model selection box (in this example set as ‘New England NSW’), c) the dialogue box providing user feedback (e.g. ‘Model New England NSW loaded’) along with a series of buttons (‘Load’, ‘Classify’, ‘Cancel’, ‘Clear’, ‘Models’) to provide the main mechanisms of user control (Fig 2).

**Figure 2:**
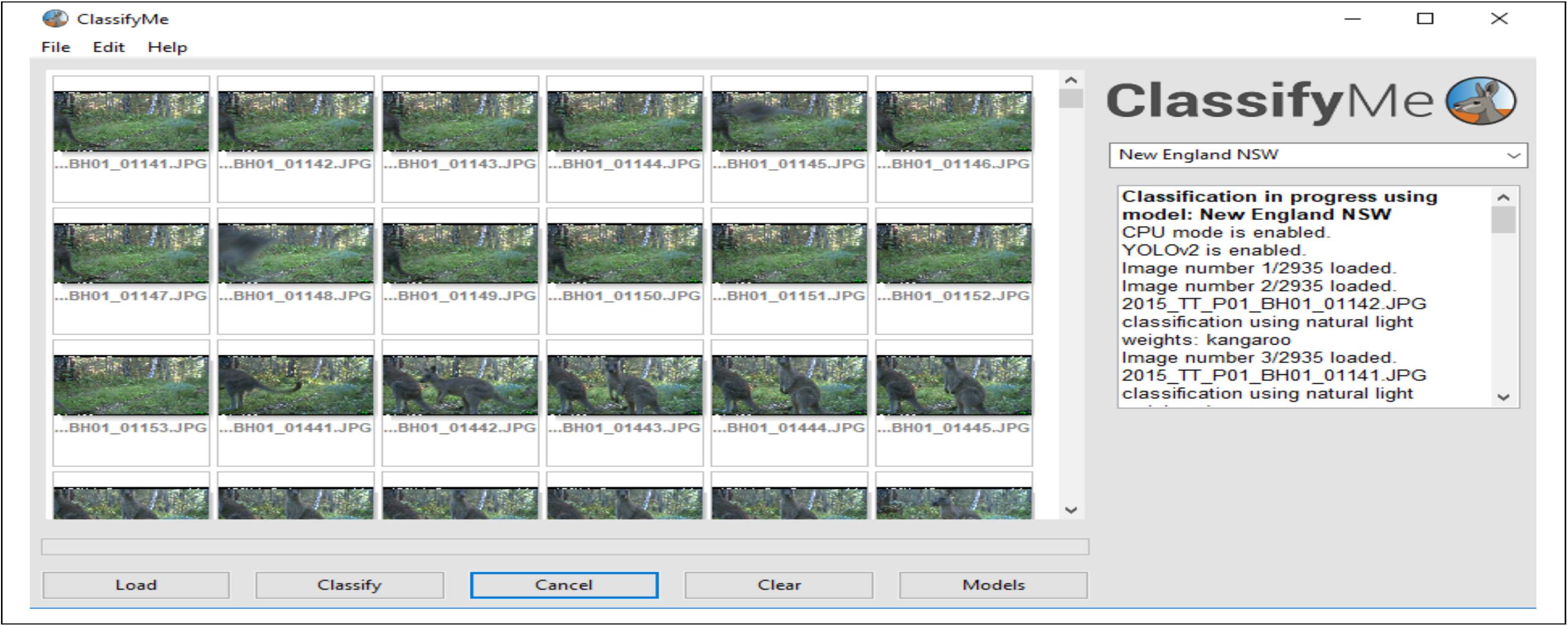
The *ClassifyMe* main user interface.

The image banner provides a useful way for the user to visually scan the contents of the image data set to confirm that the correct data set is loaded. The ‘Models’ selection box allows users to select the most appropriate detection model for their data set. *ClassifyMe* offers facilities for multiple models to be developed and offered through the web service. A user might for instance operate camera trap surveys across multiple regions (e.g. New England NSW and SW USA). Selection of a specific model allows the user to adapt the model to the specific fauna of a region. Access to specific models is dependent on user approval by the *ClassifyMe* service providers. Facilities exist for developing as many classification models as required but dependent on the provision of model training datasets.

The dialogue box of *ClassifyMe* provides the primary mechanism of user feedback with the software. It provides textual responses and prompts which guide the user through use of the software and the classification process. Finally, the GUI buttons provide the main mechanism of user control. The ‘Load’ button is used to load an image dataset from the user’s files into the system; the ‘Classify’ button to start the classification of the loaded image data using the selected model; the ‘Cancel’ button to halt the current classification task and the ‘Clear’ button to remove all current text messages from the dialogue box.

When an image dataset is loaded and the classification process started (Fig 3), each image is scanned sequentially for the presence of an animal (or other category of interest) using the selected model. *ClassifyMe* automatically sorts the images into sub-directories corresponding to the most likely classification and can also automatically detect and sort images where no animal or target category is found. The results are displayed on-screen via the dialogue box which reports the classification for each image as it is processed. The full set of classification results, which includes the confidence scores for the most likely categories, is stored as a separate csv file. *ClassifyMe* creates a separate sub-directory for each new session. The full UML structure of *ClassifyMe* (omitting security features) is described in Appendix A1.

**Figure 3:**
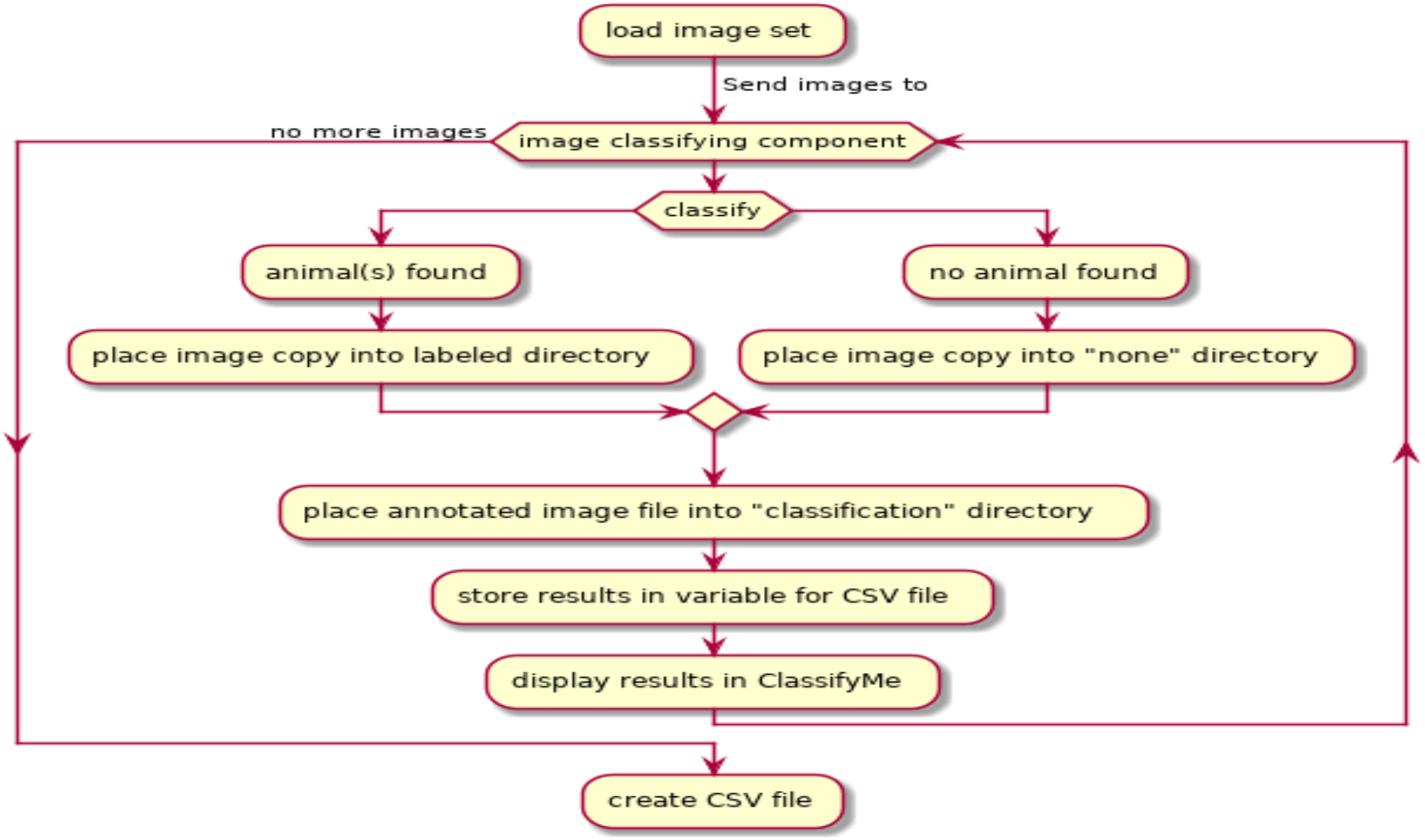
*ClassifyMe* UML diagram for image classification.

**Figure 3:**
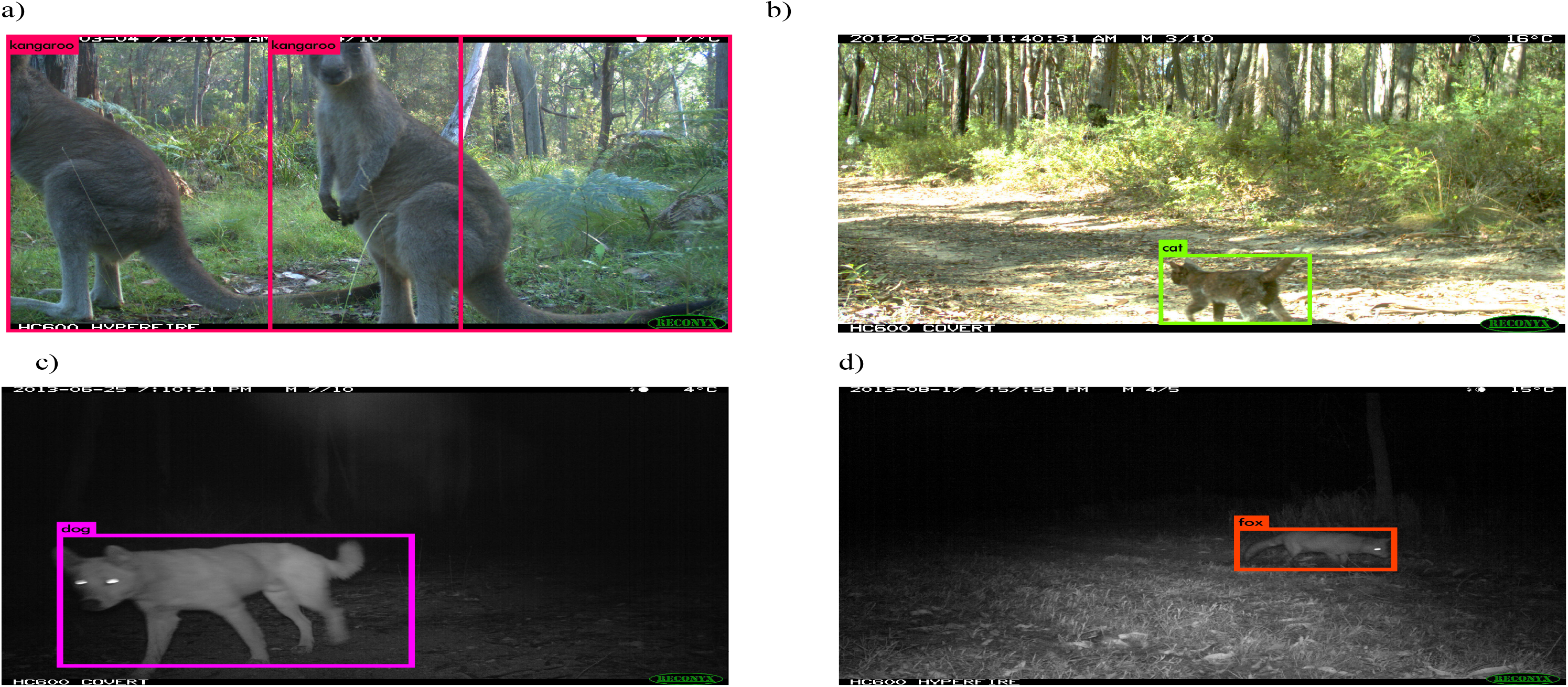
Detection Image examples from the New England dataset. (a) Macropod (Kangaroo), (b) Cat, (c) Dingo (dog) and (d) Fox.

### Recognition Models

The primary machine learning framework behind *ClassifyMe* is DarkNet and YOLOv2 (Redmon & Farhadi 2017). The YOLOv2 framework is an *object detector* deep network based on a Darknet-19 convolutional neural network structure. YOLOv2 provides access to not only a classifier (e.g. species recognition) but also a localiser (where in image) and a counter (how many animals) which facilities multi-species detections. *ClassifyMe* at present is focused on species classification but future models could incorporate these additional capabilities due to the choice of YOLOv2. YOLOv2 is designed for high-throughput processing (40-90 frames per second) whilst achieving relatively high-accuracy (YOLOv2 544 × 544 mean Average Precision 78.6@49 frames per second on Pascal VOC 2007 dataset using a GeForce GTX Titan X GPU, Redmon & Farhadi 2017). A range of other competitive object detectors such as SSD (Liu et al. 2016), Faster R-CNN (Ren et al. 2017) and R-FCN (Dai et al. 2016) could also have been selected for this task. Framework choice was governed by a range of factors including: accuracy of detection and classification; processing speed on general purpose hardware; model development and training requirements; ease of integration into other software packages and licencing. Dedicated object classifiers such as ResNet (He et al. 2016a) also provide high-accuracy performance on camera trap data (Villa, Salazar & Vargas 2017), however such models lack the future design flexibility of an object detector.

*ClassifyMe* is designed for the end-user to install relevant models from a library accessed via the configuration panel. The model is then made available for use in the model drop-down selector box e.g. the user might install the Australian and New Zealand models via the configuration panel and when analysing a specific data set select the New Zealand model. These models are developed by the *ClassifyMe* development team. Models are developed in consultation with potential end-users and when the image data provided meets the *ClassifyMe* data requirements standard (Refer Appendix A2). Importantly, *ClassifyMe* recognition models perform best when developed for the specific environment, camera trap imaging configuration and species cohort to be encountered in each study. When used outside the scope of the model, detection performance and accuracy might degrade. *ClassifyMe* is designed primarily to support end-users who have put effort into ensuring high-quality annotated datasets and who value the use of automated recognition software within their long-term study sites.

## Evaluation

*ClassifyMe* has currently been developed and evaluated for five recognition models. These are Australia (New England New South Wales), New Zealand, Serengeti (Tanzania), North America (Wisconsin) and South Western USA models. The Australia (New England NSW) dataset was developed from data collected at the University of New England’s Newholme Field Laboratory, Armidale NSW. The New Zealand model was developed as part of a predator monitoring program in the context of the *Kiwi Rescue* project (Falzon & Glen 2018). The Serengeti model was produced from a subset of the Snapshot Serengeti dataset (Swanson et al. 2015). The North America (Wisconsin) model was developed using the Snapshot Wisconsin dataset (Willi et al. 2018) whilst the South West USA was developed using data provided by Caltech camera traps data collection (Beery, Van Horn & Perona 2018). Source datasets were sub-set according to minimum data requirements for each category (comparable to the data standard advised in A2) and in light of current project developer resources.

Object detection models were developed for each dataset using YOLOv2. Hold-out test data sets were used to evaluate the performance of each model on data not used for model development. A range of model evaluation metrics were recorded including accuracy, true positive rate, positive predictive value, Matthew’s Correlation Coefficient and AUNU (Area Under the Receiver Operating Characteristic Curve of each class against the rest, using the uniform distribution) (Ferri, Hernández-Orallo & Modroiu 2009).

Overall recognition accuracies were (98.6% natural illumination, 98.7% infrared illumination) for Australia (New England, NSW), (97.9% natural and infrared illumination) for New Zealand, (99.0% natural and flash illumination) for Serengeti, (95.9% natural illumination, 98.0% infrared illumination) for North America (Wisconsin) and (96.8% natural illumination, 98.5% infrared illumination) for South West USA models. In this section, we will focus on the Australia (New England, NSW) model, further results of the other models are provided in Appendix A3.

The Australian (New England, NSW) consisted of 9 recognition classes and a total of 8,900 daylight illumination images and 8,900 infrared illumination images. Specific details of the Australian (New England, NSW) data set are provided in Table 1.

**Table 1:** Composition of New England, New South Wales, Australia data set.

| Category | Natural Sample Size<br>(Training){Validation}[Test] | IR Sample Size<br>(Training){Validation}[Test] |
| --- | --- | --- |
| Cat | (800){100}[100] | (800){100}[100] |
| Dog | (800){100}[100] | (800){100}[100] |
| Fox | (800){100}[100] | (800){100}[100] |
| Human | (800){100}[100] | (800){100}[100] |
| Macropod | (800){100}[100] | (800){100}[100] |
| Sheep | (800){100}[100] | (800){100}[100] |
| Vehicle | (800){100}[100] | (800){100}[100] |
| Other | (800){100}[100] | (800){100}[100] |
| NIL | (800){0}[100] | (800){0}[100] |

Model training was performed on a Dell XPS 8930 Intel Core i7-8700 CPU @ 3.20GHz NVIDIA GeForce GTX 1060 6GB GPU 16GB RAM 1.8TB HDD drive running a Windows 10 Professional x64 operating system using YOLOv2, via the “AlexeyAB” Windows port (https://github.com/AlexeyAB/darknet). Training consisted of 9187 epochs, 16000 iterations and 23 hours for the natural illumination model and 9820 epochs, 17000 iterations and 25 hours for the infrared illumination model.

Model performance was assessed on an independent randomly held-out test data set; the detection summary (Table 2), the confusion matrix of the specific category performance (Table 3), and the model performance metrics were evaluated (Table 4) using PyCM (Haghighi et al. 2018). Figure 4 displays examples of detection outputs, including the rectangle detection box that is overlaid on the location of the animal in the image and the detected category.

**Table 2:** Detection Summary results: New England NSW model (daylight). Randomly selected model training dataset with 800 images per class. Using threshold (Th = 0.24) to achieve a mean average precision (mAP) = 0.896067 (89.61%), 2967 detections, 993 unique truth count, and average Intersection of Union (IoU) = 75.04% and 902 True positives, 69 False Positives and 91 False Negatives. Total detection time was 20 seconds.

| Class | Average Precision |
| --- | --- |
| Cat | 99.65% |
| Dog | 90.91% |
| Fox | 90.91% |
| Human | 90.91% |
| Macropod | 80.87% |
| Sheep | 86.46% |
| Vehicle | 100.00% |
| Other | 77.14% |

**Table 3:** Confusion Matrix: New England NSW (natural illumination) model as assessed on a randomly selected hold-out test dataset.

| **Predicted** | **Actual** |  |  |  |  |  |  |  |  |  |  |
| --- | --- | --- | --- | --- | --- | --- | --- | --- | --- | --- | --- |
|  |  | Cat | Dog | Fox | Human | Macropod | NIL | Other | Sheep | Vehicle | *Precision* |
|  | Cat | 100 | 0 | 0 | 0 | 0 | 0 | 0 | 0 | 0 | 1.00 |
|  | Dog | 0 | 100 | 0 | 0 | 0 | 0 | 0 | 0 | 0 | 1.00 |
|  | Fox | 0 | 0 | 99 | 0 | 0 | 0 | 3 | 0 | 0 | 0.97 |
|  | Human | 0 | 0 | 0 | 100 | 0 | 0 | 0 | 0 | 0 | 1.00 |
|  | Macropod | 0 | 0 | 1 | 0 | 97 | 0 | 1 | 0 | 0 | 0.98 |
|  | NIL | 0 | 0 | 0 | 0 | 2 | 100 | 8 | 0 | 0 | 0.91 |
|  | Other | 0 | 0 | 0 | 0 | 0 | 0 | 91 | 0 | 0 | 1.00 |
|  | Sheep | 0 | 0 | 0 | 0 | 1 | 0 | 0 | 100 | 0 | 0.99 |
|  | Vehicle | 0 | 0 | 0 | 0 | 0 | 0 | 0 | 0 | 100 | 1.00 |
|  | *Recall* | 1.00 | 1.00 | 0.99 | 1.00 | 0.97 | 1.00 | 0.91 | 1.00 | 1.00 | **Overall Model Accuracy: 0.99** |

**Table 4:** Key Test Metrics of the New England, NSW (natural illumination) test data set. Note: * AUNU denotes Area Under Receiver Operating Characteristic Curve each class against rest using a uniform distribution.

| **Metric** | **Magnitude** |
| --- | --- |
| Overall Accuracy | 0.98556 |
| Overall Accuracy Standard Error | 0.00398 |
| 95% Confidence Interval | [0.97776,0.99335] |
| Error Rate | 0.01444 |
| Matthews Correlation Coefficient | 0.98388 |
| True Positive Rate (Macro) | 0.98556 |
| True Positive Rate (Micro) | 0.98556 |
| Positive Predictive Value (Macro) | 0.98655 |
| Positive Predictive Value (Micro) | 0.98556 |
| AUNP | 0.99187 |

The results of our testing indicate that *ClassifyMe* provides a high level of performance which is accessible across a wide range of end-user hardware with minimal configuration requirements.

## Discussion

*ClassifyMe* is the first app of its kind, it provides a software tool which allows field ecologists and wildlife managers access to the latest advances in artificial intelligence. Practitioners can utilise *ClassifyMe* to automatically identify, filter and sort camera trap image collections according to categories of interest. Such a tool fills a major gap in the operational requirements of all camera trap users irrespective of their deployments.

There are additional major benefits to localised processing on the end-user’s device. Most importantly, the local processing offered by *ClassifyMe* provides a high degree of privacy protection of end-user data. By design *ClassifyMe* does not transfer classification information of user image data back to third party, rather, all processing of the object recognition module is performed locally, with minimal user information transferred back, via encryption, to the web service. The information transferred to the web service concerns the initial registration and installation process and the on-going verification services aimed at disrupting unauthorised distribution (which is targeted specifically at poachers and similar mis-uses of *ClassifyMe* software). These privacy and data control features are known to be appealing to many in our wider network of ecological practitioners, because transmitting and sharing images with third parties compromises (1) human privacy when images contain people, (2) the location of sensitive field equipment, and (3) the location of rare and endangered species that might be targeted by illegal traffickers. Researchers and wildlife management groups also often want control over the end-use of their data and sometimes have concerns about the unforeseen consequences of unrestricted data sharing.

At present, there are few alternatives to *ClassifyMe* for the wildlife manager wanting to implement artificial intelligence technologies for the automated revision of their camera trap images. The most relevant alternative is the MLWIC: Machine Learning for Wildlife Image Classification in R package (Tabak et al. 2018). The MLWIC package provides the option to run pre-trained models and also for the user to develop their own recognition models suited to their own data sets. Whilst of benefit to a subset of research ecologists skilled with R, the approach proposed by Tabak et al. (2018) is not accessible to a wider audience as it requires investment of considerable time and effort in mastering the intricacies of the R Development Language and Environment along with the additional challenges of hardware and software configuration which are associated with this software. Integration of the MLWIC package within R is sensible if the user wants to incorporate automated image classification within their own workflows. However such automated image recognition services are already offered in other leading machine learning frameworks particularly TensorFlow (Abadi et al. 2016) and PyTorch (Paszke et al. 2017). Such frameworks offer extensive capabilities with much more memory efficient processing for a similar investment in software programming know-how (Python) and hardware configuration. In fact our wider research team routinely uses TensorFlow and PyTorch along with other frameworks such as DarkNet (Redmon & Farhadi 2017) for camera-trap focused research. Integration with R is straight-forward via exposure to a web-service API or via direct export of framework results as csv files. Within R, there are Python binding libraries which also allow access to Python code from within R and the TensorFlow interface package (Allaire & Tang 2017) also provides a comparatively easy way of accessing the full TensorFlow framework from within R. In summary, there a range of alternative options to the MLWIC package which are accessible with programming knowledge. AnimalFinder (Tack et al. 2016) is a MATLAB 2016a script available to assist with the detection of animals in time-lapse sequence camera trap images. This process is however semi-automated and does not provide species identification, it also requires access to a MATLAB software licence and corresponding software scripting skills. AnimalScanner (Yousif et al. 2019) is a similar software application providing both a MATLAB GUI and a command line executable to scan sequences of camera trap images and identify three categories (empty frames, humans or animals) based on foreground object segmentation algorithms coupled with deep learning.

The Wildlife Insights (https://wildlifeinsights.org) promises to provide cloud-based analysis services including automated species recognition but it is yet to be launched. The eMammal project provides both a cloud service and the Leopold desktop app (Forrester et al. 2013). The Leopold eMammal desktop app uses computer vision technology to search for cryptic animals within a sequence and place a bounding box around the suspected animal (He et al. 2016b). The objectives and functions of eMammal are however quite broad and support citizen science identifications, expert review, data curation and training within the context of monitoring programs and projects. This approach is very different from the approach adopted by *ClassifyMe* which is a dedicated on-demand application focused on automated species recognition on a user’s local machine with no requirement to upload datasets to 3^rd^ party sources. The iNaturalist project (https://www.inaturalist.org) is of a similar nature but focused on digital or smartphone camera acquired imagery from contributors across the world and uses deep learning convolutional neural network models to perform image recognition within its cloud platform to assist with review by citizen scientists. Whilst very useful with a wide user base, iNaturalist doesn’t specifically address the domain challenges of camera trap imagery. Motion Meerkat is a software application which also utilises computer vision in the form of mixture of Gaussian models to detect motion in videos which reduces the number of hours required for researcher review (Weinstein 2015). There is further wide range of software available including Renamer (Harris et al. 2010), Snoopy (Smedley & Terdal 2014) and VIXEN (Ramachandran & Devarajan 2017) to support camera trap data management. Young, Rode-Margono & Amin (2018) provide a detailed review of currently available camera trap software options.

An important design decision of *ClassifyMe* was not to allow end-users to train their own models. This is in contrast to software such as the MLWIC package. This decision is motivated by both legal aspects and quality control as opposed to commercial reasons. Of particular concern is use of the software to determine field locations of prized species that poachers could then target. These concerns are valid, with recent calls having been made for scientists to restrict publishing location data of highly sought-after species in peer-reviewed journals (Lindenmayer and Scheele 2017). Such capabilities could be of use to technological inclined poachers and providing such software, along with the ability to modify that software presented a number of potential legal issues. Similar concerns exist concerning human privacy legislation as outlined in Butler & Meek (2013) and Meek & Butler (2014). The strict registrations, legal and technological controls implemented within *ClassifyMe* are designed to minimise risk of misuse.

Allowing end-users to train their own models also presents quality control issues. The deep networks utilised within *ClassifyMe* (and similar software) are difficult to train to optimal performance and reliability. Specialised hardware and its configuration are also required for deep learning frameworks which can be challenging even for computer scientists. Data access and the associated labelling of datasets is also another major consideration, many users might not have sufficient sighting records nor the resources to label their datasets. The risk of developing and deploying a model which provides misleading results in practice is high with the potential consequences on wildlife observation programs quite serious. Schneider, Taylor & Kremer (2018) compared the performance of the YOLOv2 and Faster R-CNN object detectors on camera trap imagery. The YOLOv2 detector performed quite poorly with an average accuracy of 43.3 ± 14.5% (compared to Faster R-CNN which had an accuracy of 76.7 ± 8.31 %) on the Gold-Standard Snapshot Serengeti dataset. The authors suggested that the low performance was due to limited data. Our results clearly indicate that YOLOv2 can perform well with strict data quality control protocols. Furthermore, the *ClassifyMe* YOLOv2 model is most effective at longer-term study sites where the model has been calibrated using annotated data specific to the study site. *ClassifyMe* is designed to integrate well with a range of object detection frameworks including Faster-RCNN which is utilised within the software development team for research purposes. Future editions of *ClassifyMe* might also explore use of other detection frameworks or customised algorithms based on our on-going research.

*ClassifyMe* resolves the issue of model development for practitioners by out-sourcing model development to domain experts who specialise in the development of such technology in collaborative academic and government joint research programs. Users can request model development, either for private use via a commercial price or for public use which is free and on provision of image data sets to a protocol standard the model will be developed and assessed for deployment as a *ClassifyMe* model library. *ClassifyMe* is designed to enable the selection of a suitably complex model to ensure good classification performance, but to also enable storage, computation and processing within a reasonable time frame (benchmark range 1-1.5 seconds per image, Intel i7 16GB RAM) on end user computers. Cloud based solutions, such as those used in the *Kiwi Rescue* and *Wild Dog Alert* programs have the capacity to store data in a central location using a larger neural network structure on high-performance computer infrastructure. Such infrastructure is costly to run and is not ideal for all end-users.

## Conclusion

Camera trapping is commonly used to survey wildlife throughout the world but its *Achilles-heel* is the huge time and financial cost of processing data, together with the risks of human error during processing tasks. The integration of computer science and computer vision in camera trap image analysis has led to considerable advances for camera trap practitioners. The development of automated image analysis systems has filled an important gap between capturing image data in the field and analysing that data so it can be used in management decision making. *ClassifyMe* is a tool of un-matched capability for field-based operations to camera trap practitioners and organisations across the world.

## Supporting information

Model Configuration Text File

## Acknowledgements

We thank the following funding bodies for supporting our endeavours to provide a range of practitioner-based tools using current technology; Australian Wool Innovation, Meat and Livestock Australia and the Australian Government Department of Agriculture and Water Resources. This project was supported by the Centre for Invasive Animals Solutions, University of New England and the NSW Department of Primary Industries. Thanks to James Bishop, Robert Farrell, Beau Johnston, Amos Munezero, Ehsan Kiani Oshtorjani, Edmund Sadgrove, Derek Schneider, Saleh Shahinfar, Josh Stover and Jaimen Williamson for their important suggestions involving the development of the software. The data contributions from Al Glen and the Kiwi Rescue team along with Matt Gentle and Bronwyn Fancourt from Biosecurity Queensland are also greatly appreciated. Thank you to Laura Shore, Lauren Ritchie, Sally Kitto and Julie Rehwinkel of NSW DPI for their work in developing the User License Agreement.

## Appendix A1: ClassifyMe UML Structure Diagram

**Figure A1.1:**
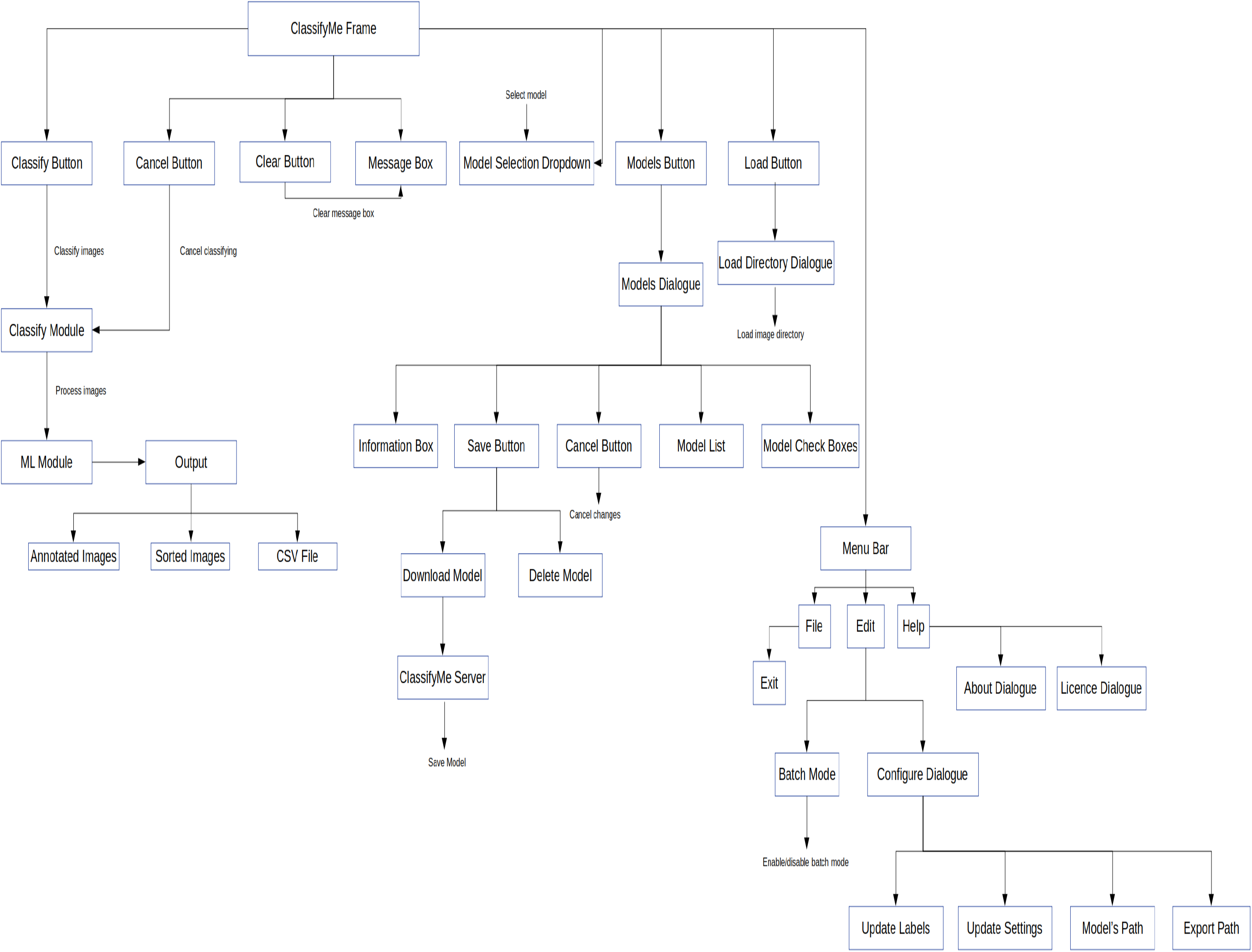
ClassifyMe Block diagram.

**Figure A1.2:**
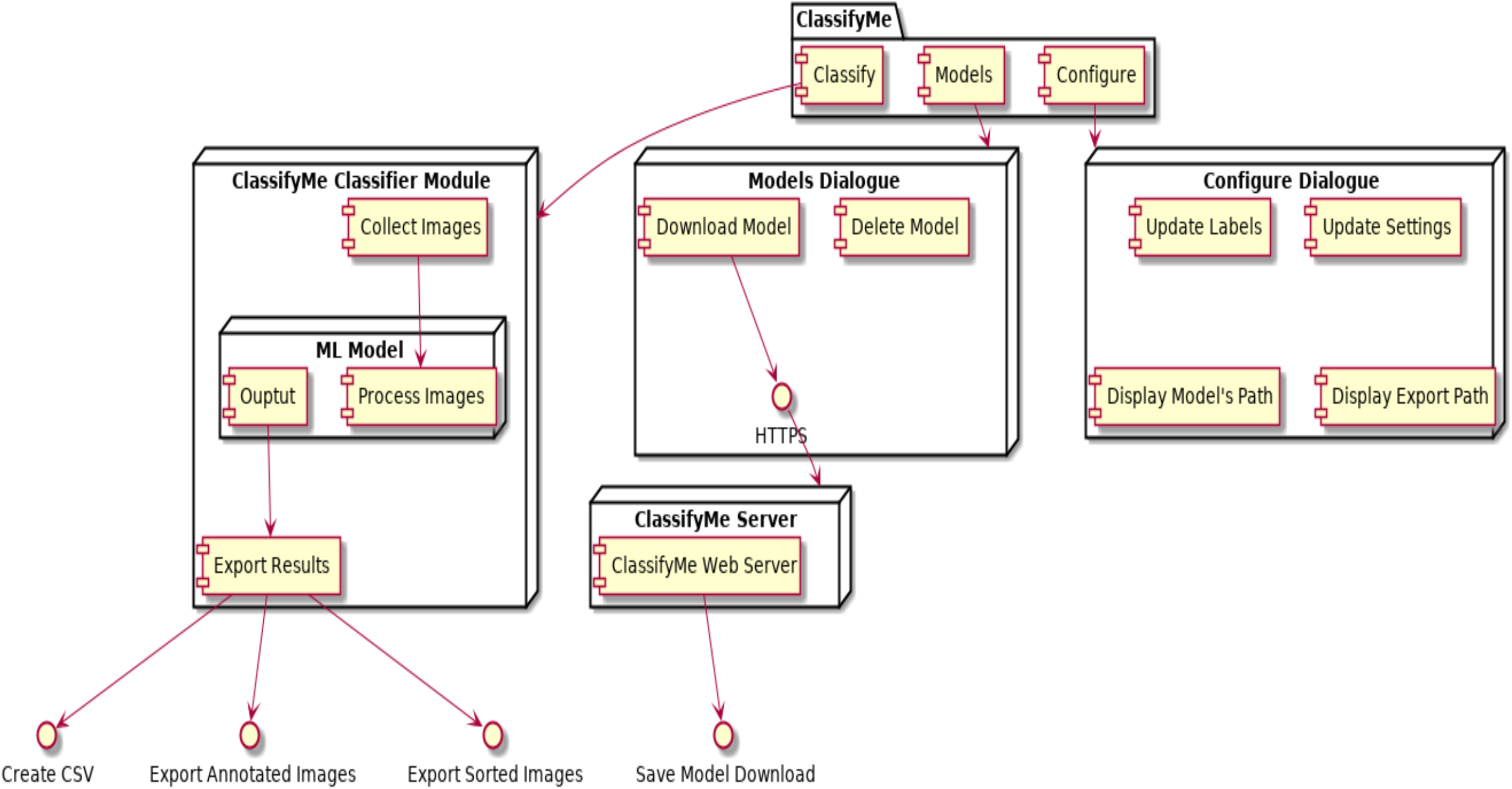
ClassifyMe Basic UML (Security Removed)

**Figure A1.3:**
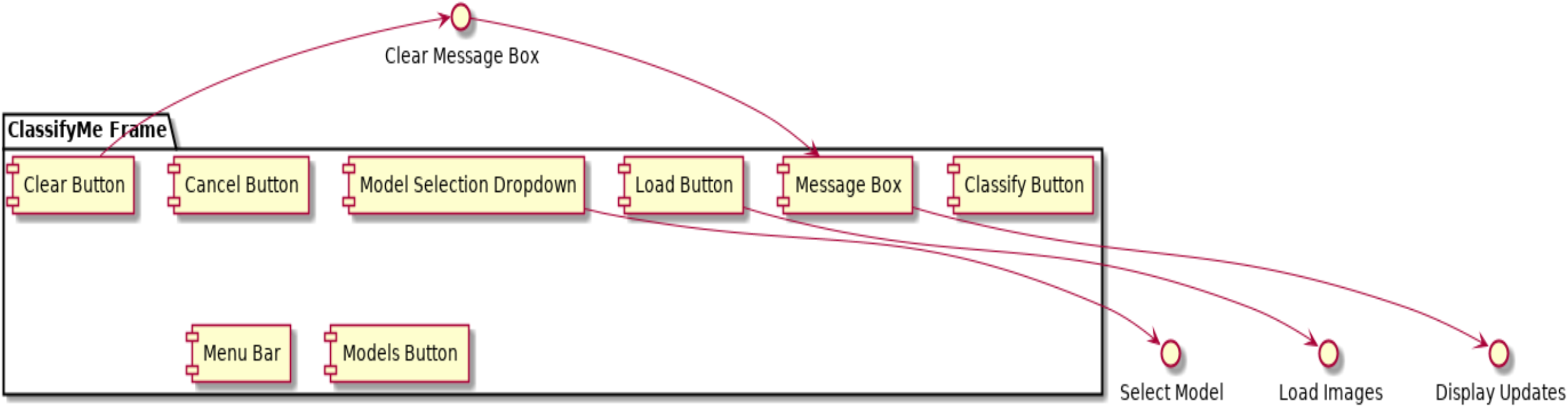
ClassifyMe Frame/GUI UML.

**Figure A1.4:**
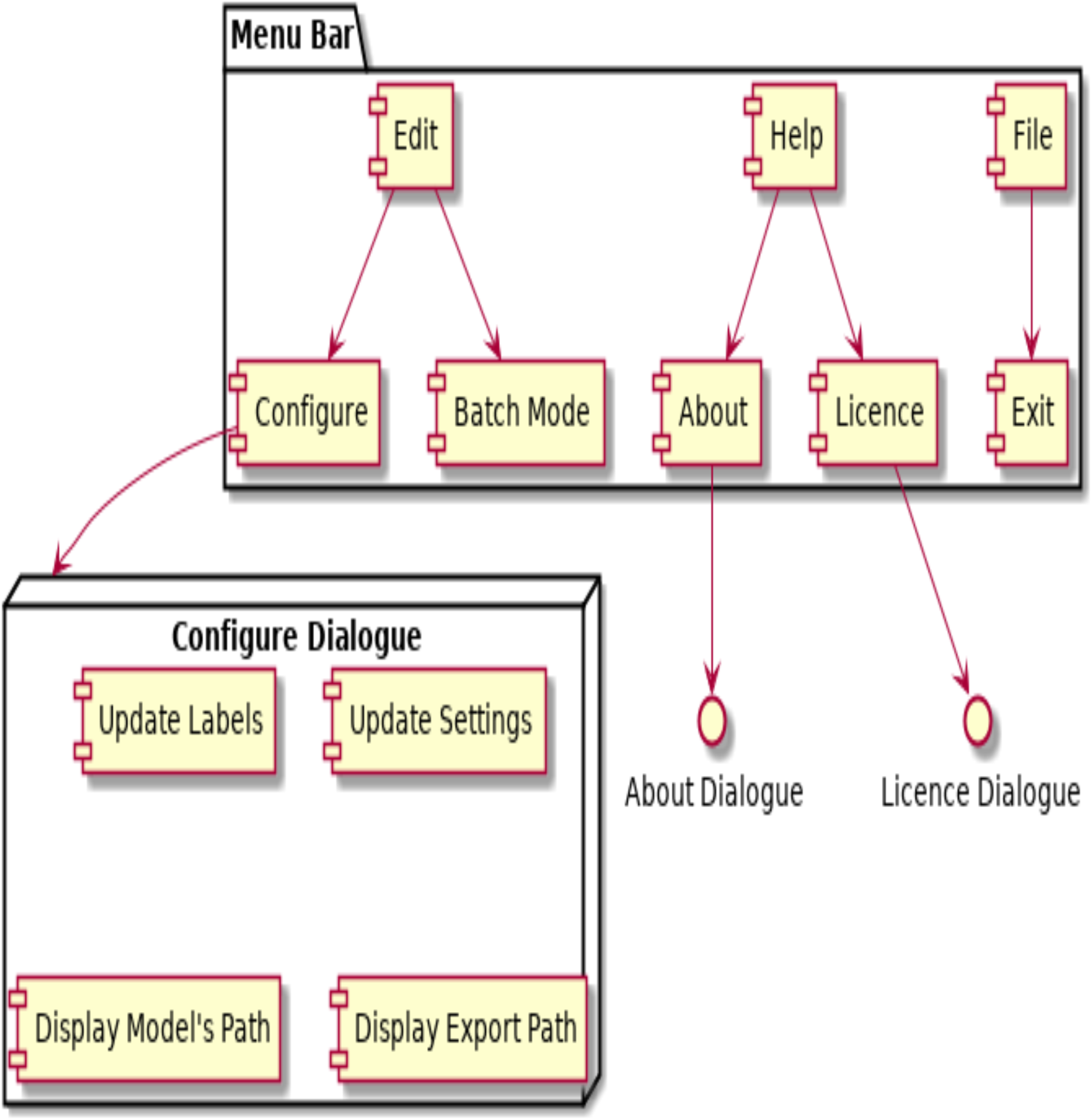
ClassifyMe Menu Bar & Configuration Dialogue.

**Figure A1.5:**
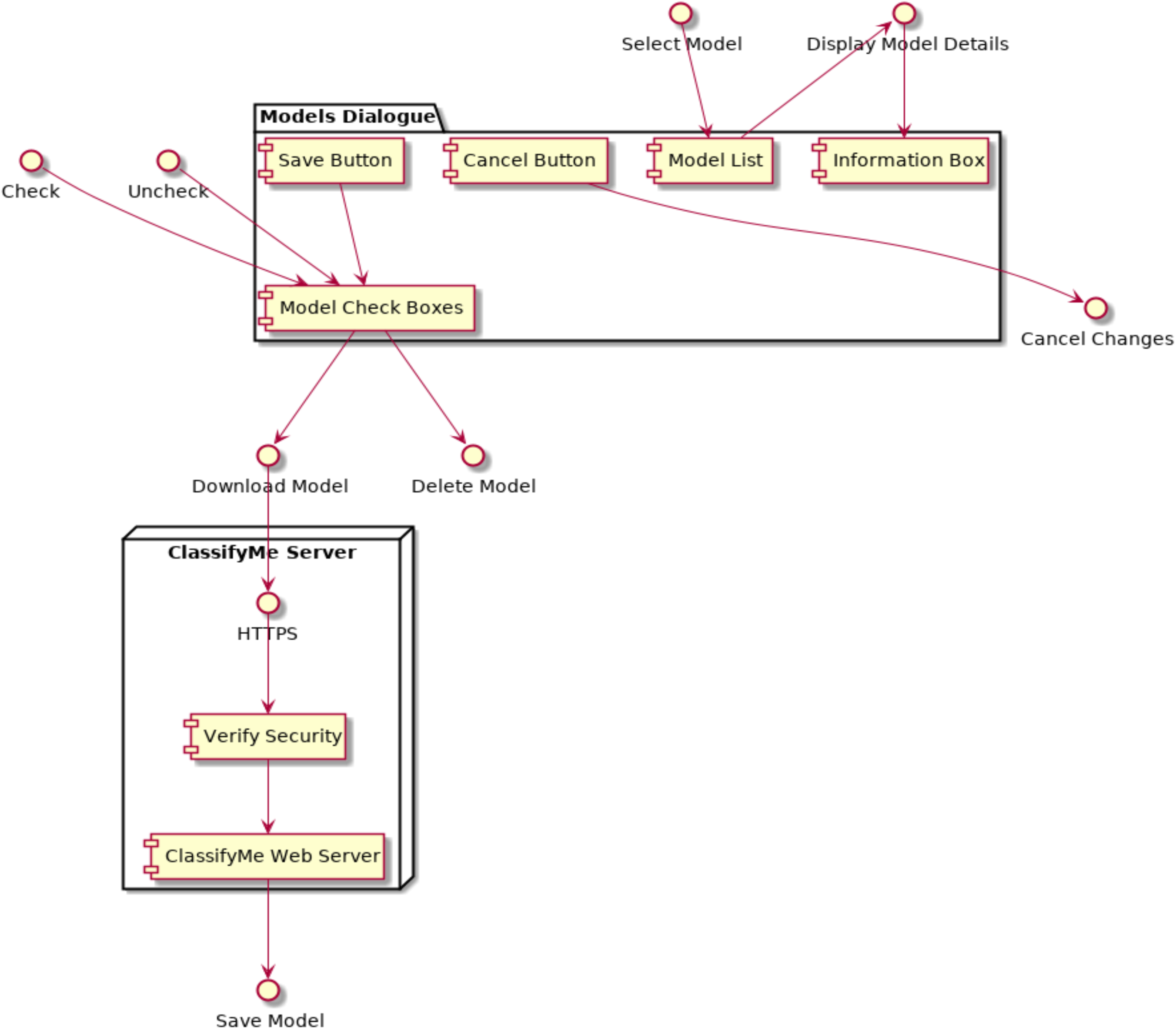
ClassifyMe Model Management.

**Figure A1.6:**
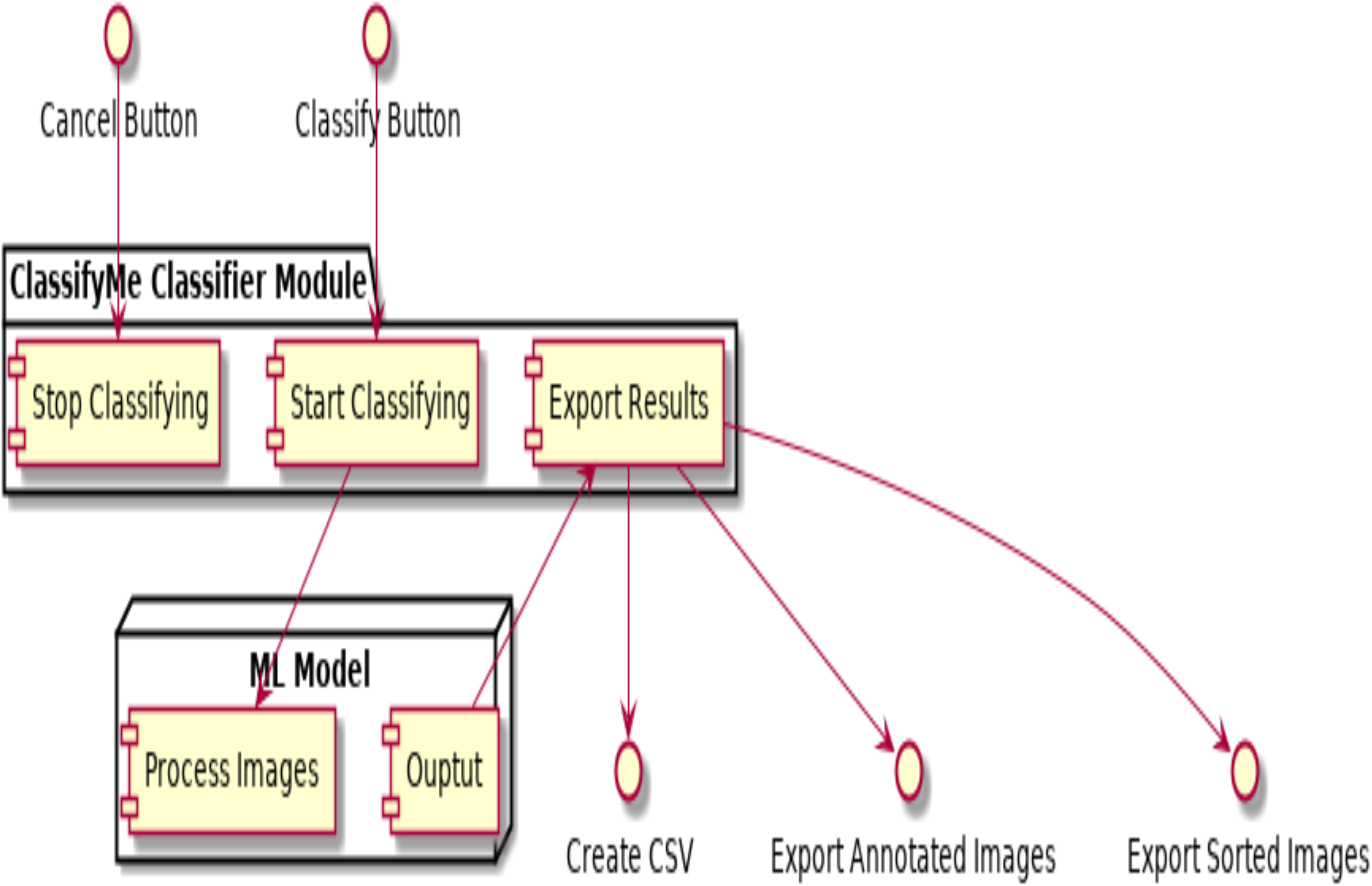
ClassifyMe Image Classification.

## Appendix A2: Data Presentation Standard for *ClassifyMe* Software

In order to develop *ClassifyMe* models for each site, our team needs the following information. Please note this form provides instructions on how we would like data to be presented for addition to the model. This document is not confirmation that UNE or NSW DPI will undertake or commit to development of a specific model. Provision of data to UNE and DPI is gratefully accepted and will not be used or provided to any other parties. Data will only be used for model development and refinement. The number of images for each category is our preferred minimum, lower numbers of images or unequal numbers of images per category or illumination class doesn’t necessarily prevent us developing a model We will assess the data provided and advise you if we cannot provide a model for your data.

### Step 1. Model Name

Please provide a suggested name for the model. This should be reflective of the location where the camera trap data was collected. Please try to limit the name to 20 characters or less e.g. CoffsCoast.

### Step 2. Metadata

Please provide the following details:

1. Principle Scientist/Senior Manager approving the provision of data
2. Institution or agency with ownership rights to data
3. Site of data collection
4. Date of collection
5. Camera trap models
6. Details of settings and placement of camera traps
7. List of the species of interest

### Step 3. Naming Convention

There is no specific naming convention, but please don’t send images with just the filename used by the camera trap. We suggest renaming files using RENAMER to a simple code that you will recognise and so that each image has a unique identifier name.

### Step 4. Model Data

We require a dictionary (folders) of all key species/category. In each folder, we require sub-folders for each illumination class e.g. Day (natural illumination), Night (IR illumination), and if used Night (*white-flash illumination). Please only organise to Genus level for species/categories. We also require false positive images which just show the landscape with no animals/objects in the frame.

These sub-folders should be further partitioned into folders describing each species/category, for example;

1. Coffs Coast/Night Infra Red/Macropod – 1230 photos
2. Coffs Coast/Day/Macropod – 1230 photos
3. Coffs Coast/ Day/False Positive – 1230 photos
4. Coffs Coast/ Night/False Positive – 1230 photos

We require equal numbers of images for each dataset i.e. each species/category per illumination class needs to have 1230 images, if you have more that is helpful but please keep the number of images consistent for each category.

### Step 5: Send Data, Forms and Information

Please contact Paul Meek to organise model development in order for us to organise our data science team resources. Data will need to be uploaded to a cloud service or sent via thumb drive or hard drive.

### Additional Notes

We currently focus on processing camera trap images where the camera has been set in the horizontal alignment and as such the camera trap PIR is horizontal with the horizon.

Please do not reduce resolution of the original camera trap images or perform any other form of image processing. Consult with us first if you have special processing requirements.

Random sampling across a variety of contexts, image backgrounds, poses, distances etc. is preferred if possible to make the system more robust.

For species/categories with insufficient data, the category ‘Other’ can be provided. This will ensure *ClassifyMe* is trained to detect animals/birds/interesting object categories in general.

Please do not send any images of humans for privacy reasons.

If you have individually recognisable animals and have enough images of them in natural and infra-red illumination, we would ask that you provide these animals in separate folder categories for example Kakudu /AnimalName/Day/Dingo and Kakudu/AnimalName/Night/Dingo. We need as many images per individual as possible but not exceeding 1000. These same animals can be included in the main data set as well.

* If you have IR and White flash photos for the night photos we need separation of IR and white flash and again 1230 images of each sub-category.

Please contact:

Dr Paul Meek

## Appendix A3: *ClassifyMe* Model Assessments

This appendix reports on the results of *ClassifyMe* applied in 5 different environments. These environments are: (i) New England Tablelands, New South Wales, Australia; (ii) Wisconsin, USA; (iii) South Western USA; (iv) Serengeti Plains, Tanzania and (v) New Zealand. Independent classification models were developed for each of these locations and where data permitted separate models within each location for natural and infrared illumination. Details of the hardware, datasets, model configuration and evaluation results are provided in order to provide a general assessment of *ClassifyMe* performance across diverse settings.

### A3.1: New England New South Wales, Australia

#### Overview

The New England region of New South Wales (NSW), Australia comprises extensive sheep stations in pastoral environments adjoining extensive sclerophyll bushland throughout the Great Dividing Range. The species cohort consists of a diverse mix of livestock, introduced and native animals. Camera trap monitoring in the region consists of either monitoring remnant vegetation on sheep stations or monitoring animal movements along vehicle access trails in state reserves and national parks. The dataset assessed represents the most frequent species or categories occurring at regional study sites for which it was possible to obtain annotations. More comprehensive sampling and species/category coverage is envisaged although the current capabilities provided by the *ClassifyMe* are already relevant and of significant assistance to research programs within this region.

**Table A3.1.1:**
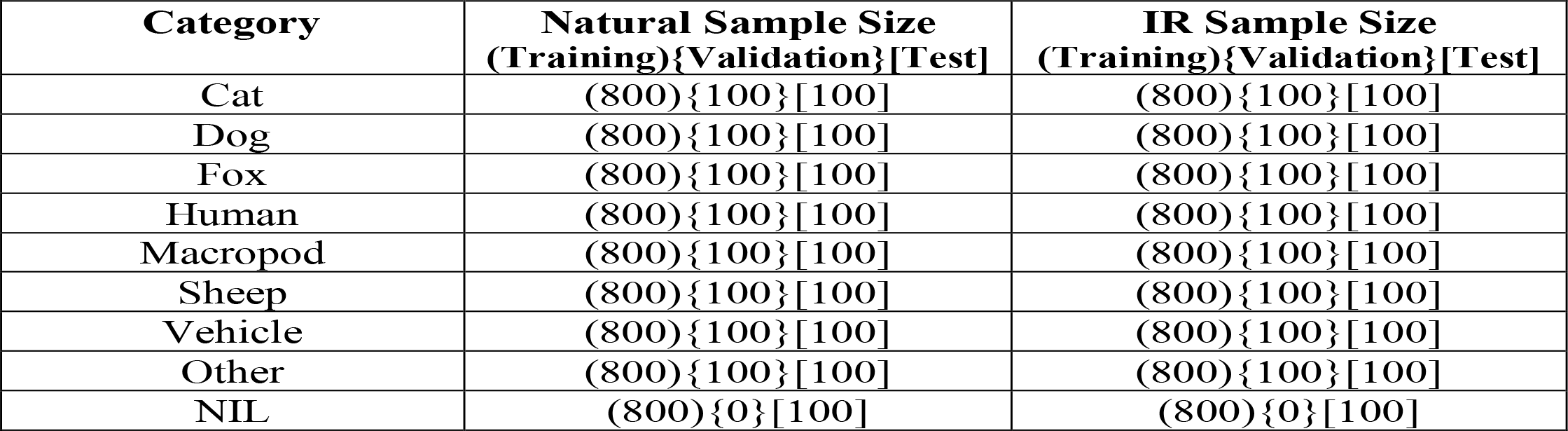
New England NSW, Australia dataset. Note the ‘Other’ dataset consisted of other animal categories of insufficient data to create a separate model category and included birds, cattle, echidnas, hares and rabbits. All images were captured using arrays of Scoutguard SG560KV or Reconyx HC600 Hyperfire cameras.

#### Model Training and Assessment Procedure

A model training dataset was formed by randomly sampling (via software) 800 images per image category in both the natural illumination and IR illumination classes. This process was done to ensure balanced numbers of samples per category in order to avoid model bias to a particular category. Separate models were developed for the natural and IR illumination classes. Data augmentation consisted of random horizontal flip, random vertical flip, random rotation of 5 degrees and random colour jitter. The YOLOv2 framework was used for model training, via the “AlexeyAB” Windows port: https://github.com/AlexeyAB/darknet. Select information is provided in Table A3.1.2 and the full YOLOv2 configuration file is provided as supplementary material (‘NewEnglandNat.cfg’) which is representative of the training procedure adopted across all models reported. Model assessment was performed by inspecting detection performance on the training dataset (Table A3.1.3), along with evaluation on the randomly held-out test datasets via confusion matrices (Tables A3.1.4 and Tables 3.1.5) and key model performance statistics (Table A3.1.6). Model assessment in the tables presented was based on *uniform* class probabilities of annotated test datasets specific to the environment. Balanced designs, with equal probability of each category allows estimation of performance metrics with no bias to a particular category.

#### Model Training Information

**Table A3.1.2:** Model Training Information, New England NSW, Australia Model.

|  | Natural Illumination | IR illumination |
| --- | --- | --- |
| **Batch Size:** | 64 | 64 |
| **Decay Rate:** | 0.0005 | 0.0005 |
| **Hours of Training:** | Approx. 31 hours | Approx. 36.5 hours |
| **Iterations (optimal weights):** | 22,700 | 26,600 |
| **Images Processed:** | 7,200 | 7,200 |
| **Learning Rate:** | 0.0001 | 0.0001 |
| **Momentum:** | 0.9 | 0.9 |

**Table A3.1.3:** Detection Summary results: New England NSW model.

| Class | Natural Illumination<br>(Average Precision) | IR Illumination<br>(Average Precision) |
| --- | --- | --- |
| Cat | 99.65% | 97.14% |
| Dog | 90.91% | 90.91% |
| Fox | 90.91% | 90.91% |
| Human | 90.91% | 90.91% |
| Macropod | 80.87% | 90.73% |
| Sheep | 86.46% | 79.93% |
| Vehicle | 100.00% | 90.91% |
| Other | 77.14% | 90.36% |

**Table A3.1.4:** Confusion Matrix: New England NSW model (natural illumination).

| Predicted | Actual |  |  |  |  |  |  |  |  |  |  |
| --- | --- | --- | --- | --- | --- | --- | --- | --- | --- | --- | --- |
|  |  | Cat | Dog | Fox | Human | Macropod | NIL | Other | Sheep | Vehicle | Precision |
|  | Cat | 100 | 0 | 0 | 0 | 0 | 0 | 0 | 0 | 0 | 1.00 |
|  | Dog | 0 | 100 | 0 | 0 | 0 | 0 | 0 | 0 | 0 | 1.00 |
|  | Fox | 0 | 0 | 99 | 0 | 0 | 0 | 3 | 0 | 0 | 0.97 |
|  | Human | 0 | 0 | 0 | 100 | 0 | 0 | 0 | 0 | 0 | 1.00 |
|  | Macropod | 0 | 0 | 1 | 0 | 97 | 0 | 1 | 0 | 0 | 0.98 |
|  | NIL | 0 | 0 | 0 | 0 | 2 | 100 | 8 | 0 | 0 | 0.91 |
|  | Other | 0 | 0 | 0 | 0 | 0 | 0 | 91 | 0 | 0 | 1.00 |
|  | Sheep | 0 | 0 | 0 | 0 | 1 | 0 | 0 | 100 | 0 | 0.99 |
|  | Vehicle | 0 | 0 | 0 | 0 | 0 | 0 | 0 | 0 | 100 | 1.00 |
|  | Recall | 1.00 | 1.00 | 0.99 | 1.00 | 0.97 | 1.00 | 0.91 | 1.00 | 1.00 | Overall Model Accuracy: 0.99 |

**Table A3.1.5:** Confusion Matrix: New England NSW model (IR illumination).

| Predicted | Actual |  |  |  |  |  |  |  |  |  |  |
| --- | --- | --- | --- | --- | --- | --- | --- | --- | --- | --- | --- |
|  |  | Cat | Dog | Fox | Human | Macropod | NIL | Other | Sheep | Vehicle | Precision |
|  | Cat | 100 | 0 | 0 | 0 | 0 | 0 | 0 | 0 | 0 | 1.00 |
|  | Dog | 0 | 100 | 0 | 0 | 0 | 0 | 0 | 0 | 0 | 1.00 |
|  | Fox | 0 | 0 | 100 | 0 | 0 | 0 | 2 | 0 | 0 | 0.98 |
|  | Human | 0 | 0 | 0 | 100 | 0 | 0 | 0 | 0 | 0 | 1.00 |
|  | Macropod | 0 | 0 | 0 | 0 | 97 | 0 | 0 | 1 | 0 | 0.99 |
|  | NIL | 0 | 0 | 0 | 0 | 1 | 100 | 6 | 0 | 0 | 0.93 |
|  | Other | 0 | 0 | 0 | 0 | 2 | 0 | 92 | 0 | 0 | 0.98 |
|  | Sheep | 0 | 0 | 0 | 0 | 0 | 0 | 0 | 99 | 0 | 1.00 |
|  | Vehicle | 0 | 0 | 0 | 0 | 0 | 0 | 0 | 0 | 100 | 1.00 |
|  | Recall | 1.00 | 1.00 | 1.00 | 1.00 | 0.97 | 1.00 | 0.92 | 0.99 | 1.00 | Overall Model Accuracy: 0.99 |

**Table A3.1.6:** Test Statistics: New England NSW.

| **Metric** | **Natural Illumination** | **IR Illumination** |
| --- | --- | --- |
| Overall Accuracy | 0.98556 | 0.98667 |
| Overall Accuracy Standard Error | 0.00398 | 0.00382 |
| 95% Confidence Interval | [0.97776,0.99335] | [0.97917,0.99416] |
| Error Rate | 0.01444 | 0.01333 |
| Matthews Correlation Coefficient | 0.98388 | 0.98506 |
| True Positive Rate (Macro) | 0.98556 | 0.98667 |
| True Positive Rate (Micro) | 0.98556 | 0.98667 |
| Positive Predictive Value (Macro) | 0.98655 | 0.98705 |
| Positive Predictive Value (Micro) | 0.98556 | 0.98667 |
| AUNP | 0.99187 | 0.9925 |

### A3.2: Wisconsin, USA

#### Overview

Wisconsin is a US State located in the MidWest and Great Lakes Region. The state features a diverse range of geographical features and environments including mixed hardwood and coniferous forests. Data was sourced from the Snapshot Wisconsin project (https://dnr.wi.gov/topic/research/projects/snapshot/) which involves a state-wide camera-trap monitoring program utilising community volunteers to annotate image data using Zooniverse (https://www.zooniverse.org) with a data subset suitable for machine learning research made available by the University of Minnesota Digital Repository (https://conservancy.umn.edu/handle/11299/199819) (Willi et al. 2018). Species and categories selected for inclusion in the *ClassifyMe* Wisconsin model are based on the frequency of images in the dataset available and guided by the data standard for development of public models as per Appendix A2. The dataset was published under the Unversity of Minnesota Data Repository *Policies and Terms of Use* (https://conservancy.umn.edu/pages/drum/policies/#terms-of-use).

**Table A3.2.1:** Wisconsin, USA dataset. Note the ‘Other’ dataset consisted of other animal categories of insufficient data to create a separate model category and included porcupine, mink, fisher, bird, bobcat, cat, otter, small mammals, opossum and sandhill crane.

| Category | Natural Sample Size<br>(Training)[Test] | IR Sample Size<br>(Training)[Test] |
| --- | --- | --- |
| Coyote | (1107) [100] | (1107) [100] |
| Deer | (1107) [100] | (1107) [100] |
| Elk | (1107) [100] | (1107) [100] |
| Hare | N/A | (1107) [100] |
| Raccoon | N/A | (1107) [100] |
| Squirrel | (1107) [100] | (1107) [100] |
| Turkey | (1107) [100] | N/A |
| Other | (1107) [100] | (1107) [100] |
| NIL | (1107)[100] | (1107)[100] |

#### Model Training and Assessment Procedure

Model training and assessment was similar to that performed in Section A3.1 for the New England, NSW model. The key differences were the source dataset, the category coverage and the number of training images per category. Tables A.3.2.1 and A.3.2.2 provide key details on the model training performance and species coverage. The ‘Other’ category incorporated random samples from the porcupine, mink, fisher, bird, bobcat, cat, otter, small mammals, opossum and sandhill crane categories. Model assessment results are reported in Tables A3.2.3, A3.2.4, A3.2.5 and A3.2.6.

#### Model Training Information

**Table A3.2.2:** Model Training Information, Wisconsin, USA Model.

|  | Natural Illumination | IR illumination |
| --- | --- | --- |
| **Batch Size:** | 64 | 64 |
| **Decay Rate:** | 0.0005 | 0.0005 |
| **Hours of Training:** | Approx. 24.8 hours | Approx. 31.3 hours |
| **Iterations (optimal weights):** | 18,000 | 22,700 |
| **Images Processed:** | 6,642 | 7,749 |
| **Learning Rate:** | 0.0001 | 0.0001 |
| **Momentum:** | 0.9 | 0.9 |

**Table A3.2.3:** Detection Summary results: Wisconsin, USA model.

| Class | Natural Illumination<br>(Average Precision) | IR Illumination<br>(Average Precision) |
| --- | --- | --- |
| Coyote | 90.80% | 90.87% |
| Deer | 90.82% | 90.77% |
| Elk | 90.89% | 90.71% |
| Hare | N/A | 90.84% |
| Raccoon | N/A | 90.87% |
| Squirrel | 90.01% | 76.72% |
| Turkey | 89.83% | N/A |
| Other | 87.97% | 90.78% |

**Table A3.2.4:** Confusion Matrix: Wisconsin, USA model (natural illumination).

| Predicted | Actual |  |  |  |  |  |  |  |  |
| --- | --- | --- | --- | --- | --- | --- | --- | --- | --- |
|  |  | Coyote | Deer | Elk | NIL | Other | Squirrel | Turkey | Precision |
|  | Coyote | 93 | 0 | 0 | 0 | 3 | 0 | 0 | 0.97 |
|  | Deer | 5 | 100 | 0 | 2 | 0 | 0 | 0 | 0.93 |
|  | Elk | 0 | 0 | 100 | 0 | 4 | 0 | 0 | 0.96 |
|  | NIL | 0 | 0 | 0 | 95 | 1 | 5 | 0 | 0.94 |
|  | Other | 1 | 0 | 0 | 1 | 91 | 1 | 0 | 0.97 |
|  | Squirrel | 0 | 0 | 0 | 1 | 0 | 92 | 0 | 0.99 |
|  | Turkey | 1 | 0 | 0 | 1 | 1 | 2 | 100 | 0.95 |
|  | Recall | 0.93 | 1.00 | 0.86 | 0.95 | 0.91 | 0.92 | 1.00 | Overall Model Accuracy: 0.96 |

**Table A3.2.5:**
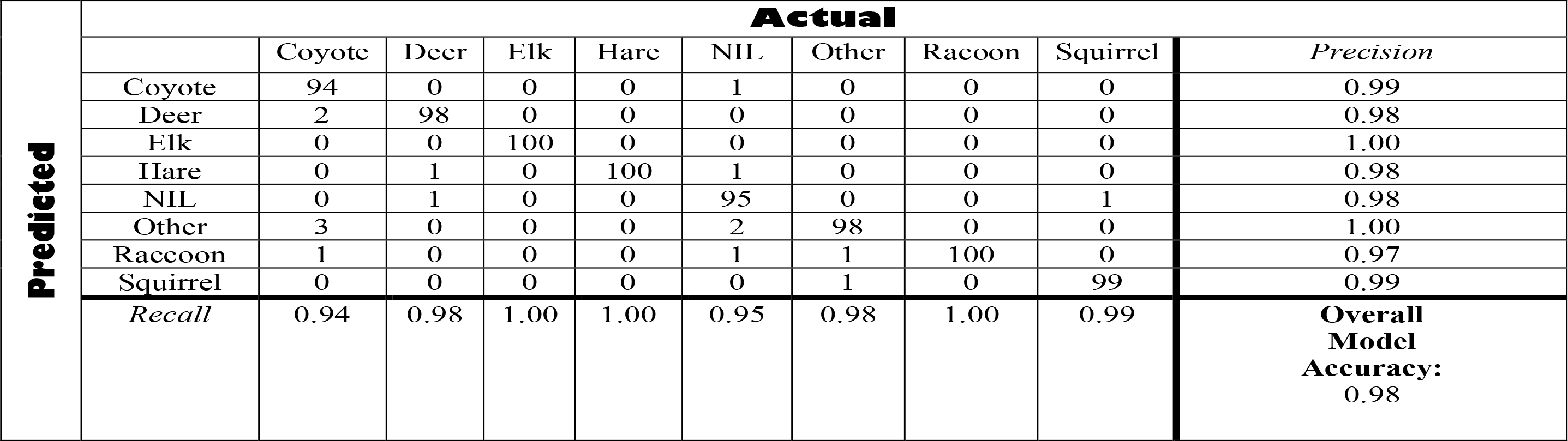
Confusion Matrix: Wisconsin, USA model (IR illumination).

**Table A3.2.6:** Test Statistics: Wisconsin, USA.

| **Metric** | **Natural Illumination** | **IR Illumination** |
| --- | --- | --- |
| Overall Accuracy | 0.95857 | 0.98 |
| Overall Accuracy Standard Error | 0.00753 | 0.00495 |
| 95% Confidence Interval | [0.94381,0.97333] | [0.9703,0.9897] |
| Error Rate | 0.04143 | 0.02 |
| Matthews Correlation Coefficient | 0.95188 | 0.97719 |
| True Positive Rate (Macro) | 0.95857 | 0.98 |
| True Positive Rate (Micro) | 0.95857 | 0.98 |
| Positive Predictive Value (Macro) | 0.95931 | 0.9802 |
| Positive Predictive Value (Micro) | 0.95857 | 0.98 |
| AUNP | 0.97583 | 0.98857 |

### A3.3: South Western, USA

#### Overview

The South Western, USA comprises camera trapping data from the Caltech Camera Traps dataset (https://beerys.github.io/CaltechCameraTraps/) with the specific data used for building the model provided by the Labeled Information Library of Alexandria: Biology and Conservation (http://lila.science/datasets/caltech-camera-traps) and has been previously reported on in Beery, Van Horn & Perona (2018). The full data set contains 243,187 images from 140 camera locations thereby representing a diverse range of environments and conditions. A random subset of the data was selected to match the *ClassifyMe* data requirements as close as possible as per Appendix A2. The Caltech camera trap dataset was published under the following Community Data Licence Agreement (https://cdla.io/permissive-1-0/).

**Table A3.3.1:** South Western, USA dataset. Note the ‘Other’ dataset consisted of other animal categories of insufficient data to create a separate model category and included skunk, deer, raccoon, opossum, car. The ‘NIL’ category was unavailable for the natural light dataset and not used in either model training or testing.

| Category | Natural Sample Size<br>(Training)[Test] | IR Sample Size<br>(Training)[Test] |
| --- | --- | --- |
| Bobcat | (1107) [100] | (1107) [100] |
| Cat | (1107) [100] | (1107) [100] |
| Coyote | (1107) [100] | (1107) [100] |
| Dog | (1107)[100] | N/A |
| Opossum | N/A | (1107) [100] |
| Other | (1107) [100] | (1107) [100] |
| Rabbit | (1107)[100] | (1107)[100] |
| Raccoon | N/A | (1107) [100] |
| Squirrel | (1107)[100] | N/A |
| NIL | (0)[100] | (0)[100] |

#### Model Training Information

**Table A3.3.2:** Model Training Information, South Western, USA Model.

|  | Natural Illumination | IR illumination |
| --- | --- | --- |
| **Batch Size:** | 64 | 64 |
| **Decay Rate:** | 0.0005 | 0.0005 |
| **Hours of Training:** | Approx. 26 hours | Approx. 26 hours |
| **Iterations (optimal weights):** | 18,000 | 18,100 |
| **Images Processed:** | 6,642 | 7,749 |
| **Learning Rate:** | 0.0001 | 0.0001 |
| **Momentum:** | 0.9 | 0.9 |

**Table A3.3.3:** Detection Summary results: South Western, USA model.

| Class | Natural Illumination<br>(Average Precision) | IR Illumination<br>(Average Precision) |
| --- | --- | --- |
| Bobcat | 90.66% | 90.75% |
| Cat | 90.81% | 90.73% |
| Coyote | 90.90% | 90.82% |
| Dog | 90.88% | N/A |
| Opossum | N/A | 90.84% |
| Other | 78.02% | 90.41% |
| Rabbit | 90.08% | 90.81% |
| Raccoon | N/A | 89.18% |
| Squirrel | 89.93% | N/A |

**Table A3.3.4:**
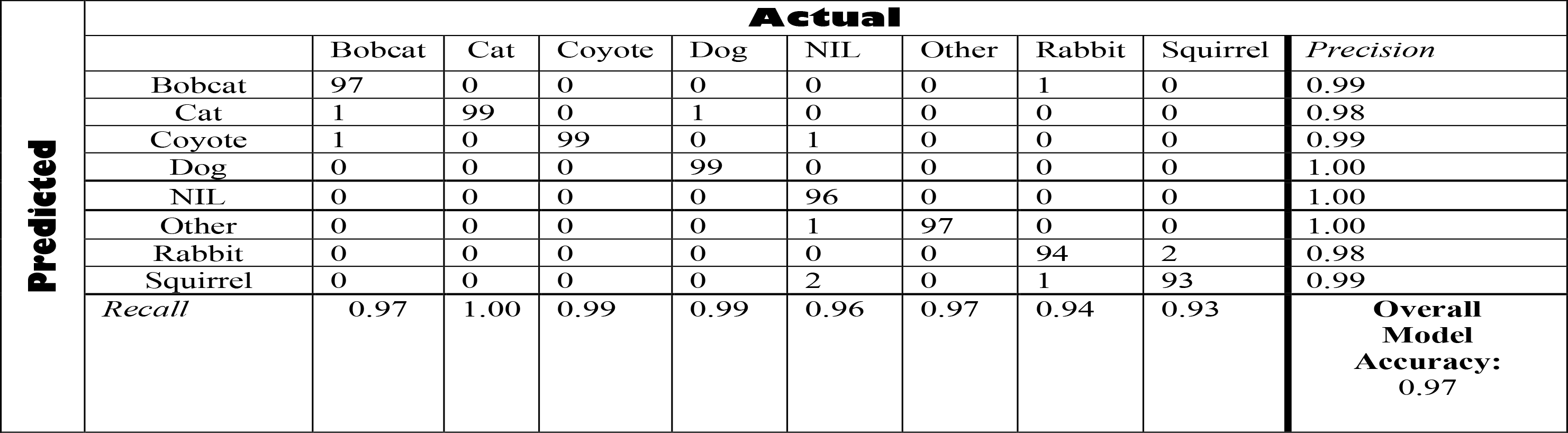
Confusion Matrix: South Western, USA model (natural illumination)

**Table A3.3.5:** Confusion Matrix: South Western, USA model (IR illumination).

| **Predicted** | **Actual** |  |  |  |  |  |  |  |  |  |
| --- | --- | --- | --- | --- | --- | --- | --- | --- | --- | --- |
|  |  | Bobcat | Cat | Coyote | Opossum | Other | NIL | Rabbit | Raccoon | *Precision* |
|  | Bobcat | 99 | 1 | 0 | 0 | 0 | 0 | 0 | 1 | 0.99 |
|  | Cat | 0 | 98 | 1 | 0 | 0 | 0 | 0 | 0 | 0.99 |
|  | Coyote | 0 | 0 | 98 | 0 | 0 | 0 | 0 | 0 | 1.00 |
|  | Opossum | 0 | 0 | 0 | 100 | 0 | 0 | 0 | 0 | 1.00 |
|  | Other | 0 | 0 | 1 | 0 | 96 | 2 | 0 | 0 | 0.97 |
|  | NIL | 0 | 1 | 0 | 0 | 0 | 98 | 0 | 0 | 0.99 |
|  | Rabbit | 1 | 0 | 0 | 0 | 2 | 0 | 100 | 0 | 0.97 |
|  | Raccoon | 0 | 0 | 0 | 0 | 0 | 0 | 0 | 99 | 1.00 |
|  | *Recall* | 0.99 | 0.98 | 0.98 | 1.00 | 0.96 | 0.98 | 1.00 | 0.99 | **Overall Model Accuracy: 0.99** |

**Table A3.3.6:** Test Statistics: South Western, USA.

| **Metric** | **Natural Illumination** | **IR Illumination** |
| --- | --- | --- |
| Overall Accuracy | 0.9675 | 0.985 |
| Overall Accuracy Standard Error | 0.00627 | 0.0043 |
| 95% Confidence Interval | [0.95521,0.97979] | [0.97658,0.99342] |
| Error Rate | 0.0325 | 0.015 |
| Matthews Correlation Coefficient | 0.96298 | 0.98287 |
| True Positive Rate (Macro) | 0.9675 | 0.985 |
| True Positive Rate (Micro) | 0.9675 | 0.985 |
| Positive Predictive Value (Macro) | 0.96883 | 0.98512 |
| Positive Predictive Value (Micro) | 0.9675 | 0.985 |
| AUNP | 0.98143 | 0.99143 |

### A3.4: Serengeti Plains, Tanzania

#### Overview

The Snapshot Serengeti project is a well-known project featuring approximately 3.2 million images and annotated by volunteers using Zooniverse (https://www.zooniverse.org). The dataset is described in Swanson et al. (2015) and features camera trap images of 40 mammal species on the African savanna. We utilised the dataset provided by the Labeled Information Library of Alexandria: Biology and Conservation (http://lila.science/datasets/snapshot-serengeti) in order to access annotated data suitable machine learning object detection. The Snapshot Serengeti dataset is highly unbalanced across categories which is reflective of the data recorded by the camera traps in operational environments. Categories selected for inclusion in the *ClassifyMe* Serengeti model were based on the most frequent categories and also meeting the minimum number of annotated images to ensure model quality control. Camera trap images included both natural and flash illumination, the *ClassifyMe* Serengeti model does not presently distinguish between these two illumination categories. Future work on the model will focus on the expansion of species and categories covered and potentially illumination specific models. This dataset was accessed under the Community Data Licence Agreement (https://cdla.io/permissive-1-0/).

**Table A3.4.1:**
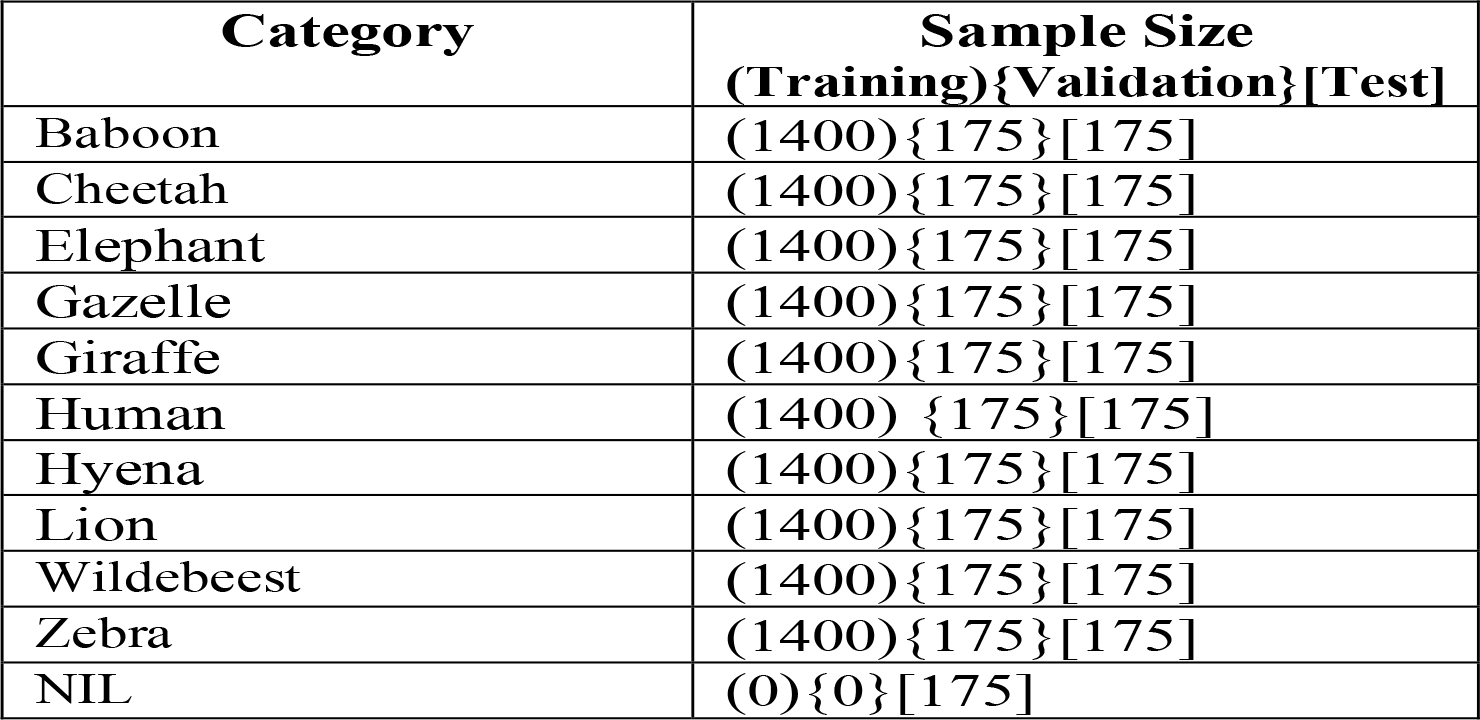
Serengeti, Tanzania dataset.

#### Model Training Information

**Table A3.4.2:** Model Training Information, Serengeti, Tanzania Model.

|  |  |
| --- | --- |
| Batch Size: | 64 |
| Decay Rate: | 0.0005 |
| Hours of Training: | Approx. 39.5 hours |
| Iterations (optimal weights): | 28,000 |
| Images Processed: | 14,000 |
| Learning Rate: | 0.0001 |
| Momentum: | 0.9 |

**Table A3.4.3:** Detection Summary results: Serengeti, Tanzania model.

| Class | Average Precision |
| --- | --- |
| Baboon | 77.43% |
| Cheetah | 88.76% |
| Elephant | 79.45% |
| Gazelle | 81.16% |
| Giraffe | 80.41% |
| Human | 90.08% |
| Hyena | 90.91% |
| Lion | 90.26% |
| Wildebeest | 79.62% |
| Zebra | 80.92% |

**Table A3.4.4:** Confusion Matrix: Serengeti, Tanzania model. WBeest denotes *Wildebeest*.

| Predicted | Actual |  |  |  |  |  |  |  |  |  |  |  |  |
| --- | --- | --- | --- | --- | --- | --- | --- | --- | --- | --- | --- | --- | --- |
|  |  | Baboon | Cheetah | Elephant | Gazelle | Giraffe | Human | Hyena | Lion | NIL | WBeest | Zebra | Precision |
|  | Baboon | 174 | 0 | 0 | 0 | 1 | 1 | 1 | 0 | 1 | 0 | 0 | 0.98 |
|  | Cheetah | 0 | 174 | 0 | 0 | 0 | 0 | 0 | 1 | 0 | 0 | 0 | 0.99 |
|  | Elephant | 0 | 0 | 175 | 0 | 0 | 1 | 0 | 0 | 0 | 0 | 0 | 0.99 |
|  | Gazelle | 0 | 0 | 0 | 172 | 0 | 0 | 0 | 0 | 0 | 0 | 0 | 1.00 |
|  | Giraffe | 0 | 0 | 0 | 0 | 174 | 0 | 0 | 0 | 3 | 0 | 0 | 0.98 |
|  | Human | 0 | 0 | 0 | 1 | 0 | 170 | 0 | 0 | 0 | 0 | 0 | 0.99 |
|  | Hyena | 0 | 0 | 0 | 0 | 0 | 0 | 174 | 1 | 0 | 0 | 0 | 0.99 |
|  | Lion | 0 | 0 | 0 | 0 | 0 | 0 | 0 | 173 | 0 | 0 | 0 | 1.00 |
|  | NIL | 1 | 0 | 0 | 2 | 0 | 3 | 0 | 0 | 170 | 0 | 0 | 0.97 |
|  | WBeest | 0 | 0 | 0 | 0 | 0 | 0 | 0 | 0 | 0 | 175 | 0 | 1.00 |
|  | Zebra | 0 | 1 | 0 | 0 | 0 | 0 | 0 | 0 | 1 | 0 | 175 | 0.99 |
|  | Recall | 0.99 | 0.99 | 1.00 | 0.98 | 0.99 | 0.97 | 0.99 | 0.99 | 0.97 | 1.00 | 1.00 | Overall Model Accuracy 0.99 |

**Table A3.4.5:** Test Statistics: Serengeti, Tanzania.

| Metric | Magnitude |
| --- | --- |
| Overall Accuracy | 0.99013 |
| Overall Accuracy Standard Error | 0.00225 |
| 95% Confidence Interval | [0.98571,0.99455] |
| Error Rate | 0.00987 |
| Matthews Correlation Coefficient | 0.98915 |
| True Positive Rate (Macro) | 0.99013 |
| True Positive Rate (Micro) | 0.99013 |
| Positive Predictive Value (Macro) | 0.99020 |
| Positive Predictive Value (Micro) | 0.99013 |
| AUNP | 0.99457 |

### A3.5: New Zealand

#### Overview

The New Zealand data set consists of camera trapping data from the *Kiwi Rescue* program (Falzon & Glen 2018). The *Kiwi Rescue* program represents an inter-institutional collaboration of government and academic scientists focused on the recovery of Kiwi (*Apteryx* spp.) populations. Camera traps were utilised in this program for predator surveillance and monitoring of Kiwi populations. Image data was obtained from sites across New Zealand, particularly Fiordland in the South Island. The dataset was provided by A. Glen of Manaaki Whenua – Landcare Research, Auckland, New Zealand and consisted of 5,228 annotated camera trap images (9.57GB) relevant to New Zealand Kiwi population monitoring. Due to the limited number of images per category across illumination classes, the data set was combined into monochrome format by transforming RGB colour images to grey-scale images. Therefore, only a single model was developed and evaluated for the NZ dataset.

**Table A3.5.1:** New Zealand dataset.

| Category | Sample Size<br>(Training){Validation}[Test] |
| --- | --- |
| Bird | (714){94}[20] |
| Cat | (174){21}[20] |
| Hedgehog | (765){101}[20] |
| Kiwi | (837){110}[20] |
| Sheep | (940){125}[20] |
| Stoat | (502){66}[20] |
| NIL | (523){65}[20] |

#### Model Training Information

**Table A3.5.2:** Model Training Information, New Zealand Model.

|  |  |
| --- | --- |
| Batch Size: | 64 |
| Decay Rate: | 0.0005 |
| Hours of Training: | Approx. 24.2 hours |
| Iterations (optimal weights): | 19,400 |
| Images Processed: | 4,455 |
| Learning Rate: | 0.0001 |
| Momentum: | 0.9 |

**Table A3.5.3:** Detection Summary results: New Zealand Model.

| Class | Average Precision |
| --- | --- |
| Bird | 80.61% |
| Cat | 90.91% |
| Hedgehog | 90.91% |
| Kiwi | 90.91% |
| Sheep | 90.43% |
| Stoat | 90.91% |

**Table A3.5.4:**
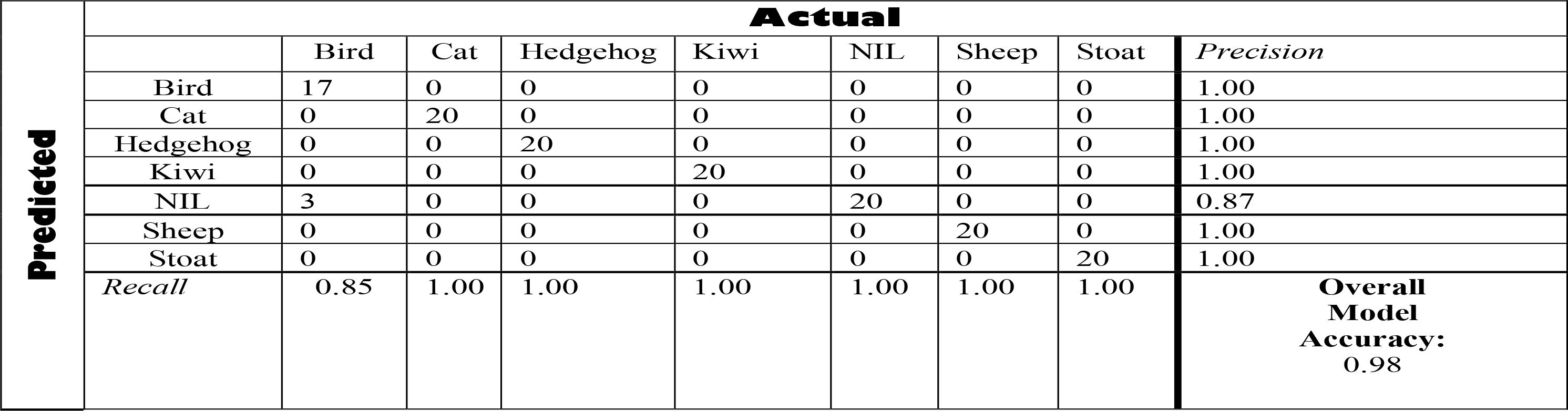
Confusion Matrix: New Zealand Model.

**Table A3.5.5:** Test Statistics: New Zealand.

| Metric | Magnitude |
| --- | --- |
| Overall Accuracy | 0.97857 |
| Overall Accuracy Standard Error | 0.01224 |
| 95% Confidence Interval | [0.95458,1.00256] |
| Error Rate | 0.02143 |
| Matthews Correlation Coefficient | 0.97552 |
| True Positive Rate (Macro) | 0.97857 |
| True Positive Rate (Micro) | 0.97857 |
| Positive Predictive Value (Macro) | 0.98137 |
| Positive Predictive Value (Micro) | 0.97857 |
| AUNP | 0.9875 |

### A3.6: Computational Performance Evaluations

**Table A3.6.1:**
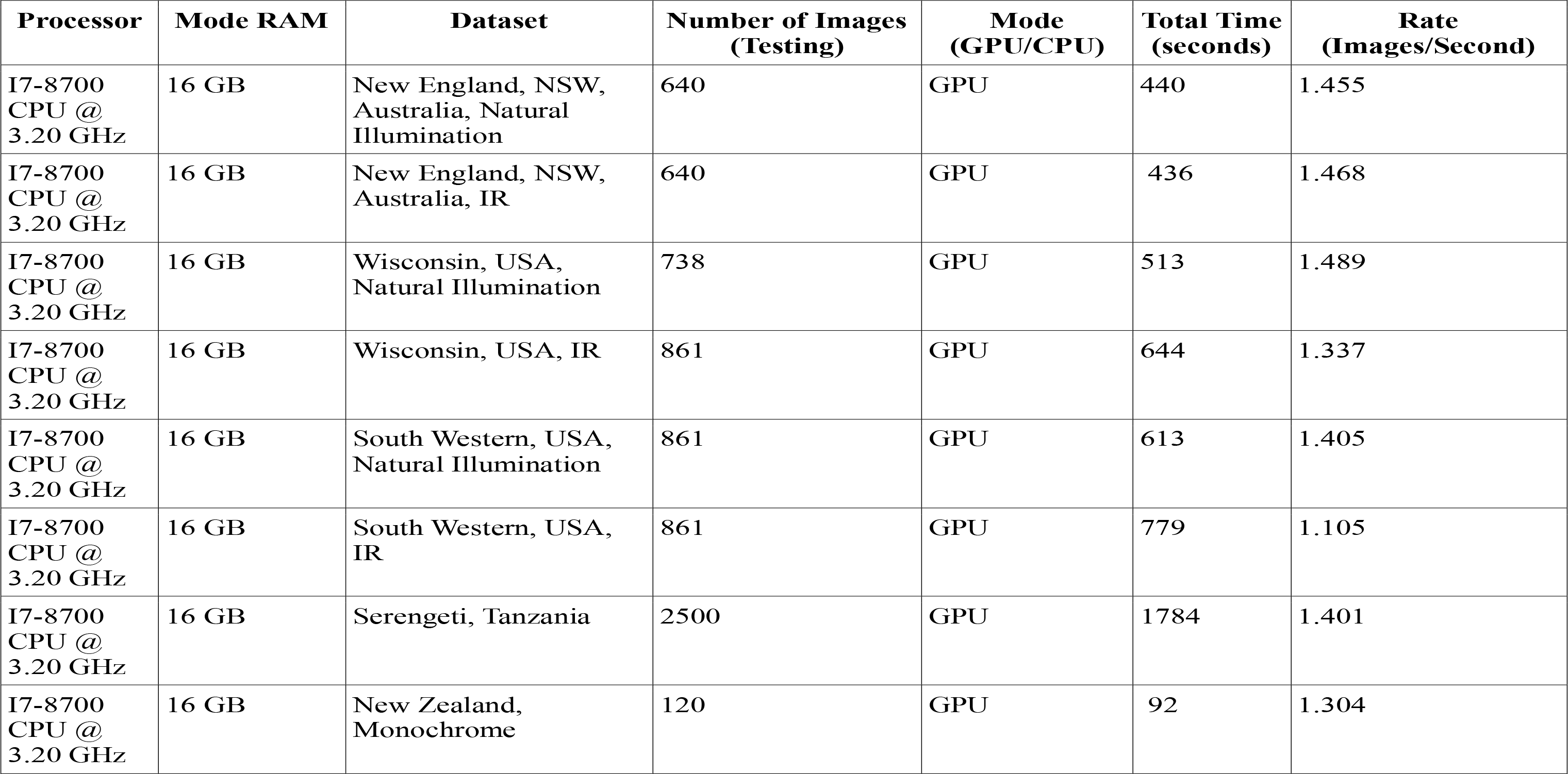
ClassifyMe computational performance evaluation metrics.

| Processor | Mode RAM | Dataset | Number of Images (Testing) | Mode (GPU/CPU) | Total Time (seconds) | Rate (Images/Second) |
| --- | --- | --- | --- | --- | --- | --- |
| I7-8700<br>CPU @<br>3.20 GHz | 16 GB | New England, NSW, Australia, Natural Illumination | 640 | GPU | 440 | 1.455 |
| I7-8700<br>CPU @<br>3.20 GHz | 16 GB | New England, NSW, Australia, IR | 640 | GPU | 436 | 1.468 |
| I7-8700<br>CPU @<br>3.20 GHz | 16 GB | Wisconsin, USA, Natural Illumination | 738 | GPU | 513 | 1.489 |
| I7-8700<br>CPU @<br>3.20 GHz | 16 GB | Wisconsin, USA, IR | 861 | GPU | 644 | 1.337 |
| I7-8700<br>CPU @<br>3.20 GHz | 16 GB | South Western, USA, Natural Illumination | 861 | GPU | 613 | 1.405 |
| I7-8700<br>CPU @<br>3.20 GHz | 16 GB | South Western, USA, IR | 861 | GPU | 779 | 1.105 |
| I7-8700<br>CPU @<br>3.20 GHz | 16 GB | Serengeti, Tanzania | 2500 | GPU | 1784 | 1.401 |
| I7-8700<br>CPU @<br>3.20 GHz | 16 GB | New Zealand, Monochrome | 120 | GPU | 92 | 1.304 |

## References

Abadi, M., Barham, P., Chen, J., Chen, Z., Davis, A., Dean, J., Devin, M., Ghemawat, S., Irving, G., Isard, M. & Kudlur, M. (2016, November). TensorFlow: a system for large-scale machine learning. In OSDI (Vol. 16, pp. 265–283).

Allaire, J.J. & Tang Y. (2017) TensorFlow: R Interface to ‘TensorFlow’. R package version 1.4.3. https://tensorflow.rstudio.com (Retrieved 22 May 2019).

Ahumada, J.A., Silva, C.E., Gajapersad, K., Hallam, C., Hurtado, J., Martin, E., McWilliam, A., Mugerwa, B., O’Brien, T., Rovero, F., Sheil, D., Spironello, W.R., Winarni, N., & Andelman, S.J. (2011) Community structure and diversity of tropical forest mammals: data from a global camera trap network. Philosophical Transactions of the Royal Society B: Biological Sciences, 366, 2703–11.

Beery, S., Van Horn, G., & Perona, P. (2018). Recognition in Terra Incognita. In Proceedings of the European Conference on Computer Vision (ECCV) (pp. 456–473).

Bennett, E. L. (2015). Legal ivory trade in a corrupt world and its impact on African elephant populations. Conservation Biology, 29(1), 54–60.

Butler, D. A., & Meek, P. (2013). Camera trapping and invasions of privacy: an Australian legal perspective. Torts Law Journal, 20(3), 235–264.

Claridge, A.W. and Paull, D.J. (2014) How long is a piece of string? Camera trapping methodology is question dependent. In: Camera Trapping: Wildlife Management and Research. (Ed. P.D., Meek, Banks, P. B., Claridge, A. W., Fleming, P. J. S., Sanderson, J. G., and Swann, D.) (pp. 205–214). (CSIRO Publishing: Melbourne, Australia).

Csillik, O., Cherbini, J., Johnson, R., Lyons, A., & Kelly, M. (2018). Identification of Citrus Trees from Unmanned Aerial Vehicle Imagery Using Convolutional Neural Networks. Drones, 2(4), 39.

Dai, J., Li, Y., He, K., & Sun, J. (2016). R-fcn: Object detection via region-based fully convolutional networks. In Advances in Neural Information Processing Systems (pp. 379–387).

Esteva, A., Kuprel, B., Novoa, R. A., Ko, J., Swetter, S. M., Blau, H. M., & Thrun, S. (2017). Dermatologist-level classification of skin cancer with deep neural networks. Nature, 542(7639), 115.

Falzon, G., Meek, P.D., & Vernes, K. (2014). Computer-assisted identification of small Australian mammals in camera trap imagery. In: Camera Trapping: Wildlife Management and Research. (Ed. P.D., Meek, Banks, P. B., Claridge, A. W., Fleming, P. J. S., Sanderson, J. G., and Swann, D.) (pp. 299–306). (CSIRO Publishing: Melbourne, Australia).

Falzon, G., & Glen, A. (2018). Developing image recognition software for New Zealand animals. In: Proceedings of the 31st Australasian Wildlife Management Society Conference. (p. 102). (Australasian Wildlife Management Society, Hobart, Australia).

Ferri, C., Hernández-Orallo, J., & Modroiu, R. (2009). An experimental comparison of performance measures for classification. Pattern Recognition Letters, 30(1), 27–38.

Forrester, T., McShea, W. J., Keys, R. W., Costello, R., Baker, M., & Parsons, A. (2013). eMammal–citizen science camera trapping as a solution for broad-scale, long-term monitoring of wildlife populations. Sustainable Pathways: Learning from the Past and Shaping the Future.

Gormley, A. M., Forsyth, D. M., Griffioen, P., Lindeman, M., Ramsey, D. S., Scroggie, M. P., & Woodford, L. (2011). Using presence-only and presence–absence data to estimate the current and potential distributions of established invasive species. Journal of Applied Ecology, 48(1), 25–34.

Gowen, C., & Vernes, K. (2014). Population estimates of an endangered rock wallaby, *Petrogale penicillata,* using time-lapse photography. In: Camera Trapping: Wildlife Management and Research. (Ed. P.D., Meek, Banks, P. B., Claridge, A. W., Fleming, P. J. S., Sanderson, J. G., and Swann, D.) (pp. 61–68). (CSIRO Publishing: Melbourne, Australia).

Haghighi, S., Jasemi, M., Hessabi, S., & Zolanvari, A. (2018). PyCM: Multiclass confusion matrix in Python. The Journal of Open Source Software, 3(25), 729.

Harmsen, B. J., Foster, R. J., Silver, S. C., Ostro, L. E., & Doncaster, C. P. (2009). Spatial and temporal interactions of sympatric jaguars (*Panthera onca*) and pumas (*Puma concolor*) in a neotropical forest. Journal of Mammalogy, 90(3), 612–620.

Harris, G., Thompson, R., Childs, J. L., & Sanderson, J. G. (2010). Automatic storage and analysis of camera trap data. The Bulletin of the Ecological Society of America, 91(3), 352–360.

He, K., Zhang, X., Ren, S., & Sun, J. (2016a). Deep residual learning for image recognition. In Proceedings of the IEEE Conference on Computer Vision and Pattern Recognition (pp. 770–778).

He, Z., Kays, R., Zhang, Z., Ning, G., Huang, C., Han, T. X., Millspaugh, J., Forrester, T. & McShea, W. (2016b). Visual informatics tools for supporting large-scale collaborative wildlife monitoring with citizen scientists. IEEE Circuits and Systems Magazine, 16(1), 73–86.

Jackson, R. M., Roe, J. D., Wangchuk, R., & Hunter, D.O. (2006). Estimating snow leopard population abundance using photography and capture-recapture techniques. Wildlife Society Bulletin, 34(3), 772–781.

Karanth, K. U. (1995). Estimating tiger *Panthera tigris* populations from camera-trap data using capture-recapture models. Biological Conservation, 71(3), 333–338.

Khorozyan, I. G., Malkhasyan, A. G., & Abramov, A. V. (2008). Presence–absence surveys of prey and their use in predicting leopard (*Panthera pardus*) densities: a case study from Armenia. Integrative Zoology, 3(4), 322–332.

Lindenmayer, D., & Scheele, B. (2017). Do not publish. Science 356 (6340), 800–801.

Linkie, M., & Ridout, M. S. (2011). Assessing tiger–prey interactions in Sumatran rainforests. Journal of Zoology, 284(3), 224–229.

Liu, W., Anguelov, D., Erhan, D., Szegedy, C., Reed, S., Fu, C. Y., & Berg, A. C. (2016). SSD: Single shot multibox detector. In: Leibe B., Matas J., Sebe N., Welling M. (eds) Computer Vision –ECCV 2016. ECCV 2016. Lecture Notes in Computer Science, vol 9905 (pp. 21–37). (Springer, Cham).

Meek, P. D., Zewe, F., & Falzon, G. (2012). Temporal activity patterns of the swamp rat (*Rattus lutreolus*) and other rodents in north-eastern New South Wales, Australia. Australian Mammalogy, 34(2), 223–233.

Meek, P. D., Vernes, K., & Falzon, G. (2013). On the reliability of expert identification of small-medium sized mammals from camera trap photos. Wildlife Biology in Practice, 9(2), 1–19.

Meek, P. D., & Butler, D. (2014). Now we can ‘see the forest and the trees too’ but there are risks: camera trapping and privacy law in Australia. In: Camera Trapping: Wildlife Management and Research. (Ed. P.D., Meek, Banks, P. B., Claridge, A. W., Fleming, P. J. S., Sanderson, J. G., and Swann, D.) (pp. 331–345). (CSIRO Publishing: Melbourne, Australia).

Meek, P.D., Ballard, A.G., Banks, P.B., Claridge, A.W., Fleming, P.J.S., Sanderson, J.G, Swann, D.E. (Eds) (2014a). ’Camera Trapping in Wildlife Research and Monitoring.’ (CSIRO Publishing: Melbourne, Australia)

Meek, P.D. Fleming, P.J.S., Ballard, G., Banks, P., Claridge, A.W., McMahon, S., Sanderson J. & Swann, D.E. (2014b). Putting contemporary camera trapping in focus. In ’Camera Trapping in Wildlife Research and Management.’ (Eds PD Meek, AG Ballard, PB Banks, AW Claridge, PJS Fleming, JG Sanderson, DE Swann.) (pp 349–356). (CSIRO Publishing: Melbourne, Australia).

Meek, P.D., Ballard, G.A., Falzon, G., Williamson, J., Milne, H., Farrell, R., Stover, J., Mather-Zardain, A. T., Bishop, J., C. Ka-Wai Cheung, E., Lawson, C. K., Munezero, A. M., Schneider, D., Johnston, B. E., Kiani, E., Shahinfar, S., Sadgrove, E. J., & Fleming, P.J.S. (in press). Camera Trapping Technology and Advances: into the New Millennium. Australian Zoologist Special Edition: Phones, drones and people: new approaches to zoology.

Norouzzadeh, M. S., Nguyen, A., Kosmala, M., Swanson, A., Palmer, M. S., Packer, C., & Clune, J. (2018). Automatically identifying, counting, and describing wild animals in cameratrap images with deep learning. Proceedings of the National Academy of Sciences, 201719367.

O’Connell, A.F., Nichols, J.D., & Karanth, K.U. (2011) Camera Traps in Animal Ecology Methods and Analyses. Springer: New York.

Paszke, A., Gross, S., Chintala, S., Chanan, G., Yang, E., DeVito, Z., Desmaison, A., Antiga, L., & Lerer, A. (2017). Automatic differentiation in PyTorch.

Ramachandran, P., & Devarajan, K. (2018). ViXeN: An open-source package for managing multimedia data. Methods in Ecology and Evolution, 9(3), 785–792.

Ramsey, D. S., Caley, P. A., & Robley, A. (2015). Estimating population density from presence–absence data using a spatially explicit model. The Journal of Wildlife Management, 79(3), 491–499.

Redmon, J., & Farhadi, A. (2017). YOLO9000: better, faster, stronger. In The IEEE Conference on Computer Vision and Pattern Recognition (CVPR) (pp. 7263–7271). (IEEE).

Ren, S., He, K., Girshick, R., & Sun, J. (2017). Faster R-CNN: towards real-time object detection with region proposal networks. IEEE Transactions on Pattern Analysis & Machine Intelligence, 6:1137–1149.

Rovero, F. & Zimmerman, F. (eds). (2016). Camera Trapping for Wildlife Research. Exeter: Pelagic Publishing UK.

Schneider, S., Taylor, G. W., & Kremer, S. C. (2018). Deep Learning Object Detection Methods for Ecological Camera Trap Data. In 2018 15th Conference on Computer and Robot Vision (CRV) (pp. 321–328). (IEEE).

Smedley, R. & Terdal, E. (2014). Snoopy: portable software for capture-recapture surveys. Conference paper.

Swann, D.E. & Perkins, N. (2014) Camera trapping for animal monitoring and management: a review of applications. In ‘Camera Trapping in Wildlife Research and Management.’ In: Camera Trapping: Wildlife Management and Research. (Ed. P.D., Meek, Banks, P. B., Claridge, A. W., Fleming, P. J. S., Sanderson, J. G., and Swann, D.) (pp. 4–11). (CSIRO Publishing: Melbourne, Australia).

Swanson, A., Kosmala, M., Lintott, C., Simpson, R., Smith, A., & Packer, C. (2015). Snapshot Serengeti, high-frequency annotated camera trap images of 40 mammalian species in an African savanna. Scientific Data, 2, 150026.

Tabak, M. A., Norouzzadeh, M. S., Wolfson, D. W., Sweeney, S. J., VerCauteren, K. C., Snow, N. P., Halseth, J.M., Di Salvo, P.A., Lewis, J.S., White, M.D., & Teton, B. (2018). Machine learning to classify animal species in camera trap images: applications in ecology. Methods in Ecology and Evolution, 10(4): 585–590.

Tack, J. L. P., West, B. S., McGowan, C. P., Ditchkoff, S. S., Reeves, S. J., Keever, A. C., & Grand, J. B. (2016). AnimalFinder: A semi-automated system for animal detection in timelapse camera trap images. Ecological Informatics, 36, 145–151.

Trolle, M., & Kéry, M. (2003). Estimation of ocelot density in the Pantanal using capturerecapture analysis of camera-trapping data. Journal of Mammalogy, 84(2), 607–614.

Vernes, K., Smith, M., & Jarman, P. (2014). A novel camera-based approach to understanding the foraging behaviour of mycophagous mammals. In: Camera Trapping: Wildlife Management and Research. (Ed. P.D., Meek, Banks, P. B., Claridge, A. W., Fleming, P. J. S., Sanderson, J. G., and Swann, D.) (pp. 215–224). (CSIRO Publishing: Melbourne, Australia).

Vernes, K., & Jarman, P. (2014). Foraging behaviour and handling times for long-nosed potoroos (*Potorous tridactylus*) foraging for buried truffles. Australian Mammalogy 36:128–130.

Vernes, K., Sangay, T., Rajaratnam, R., & Singye, R. (2015). Social interaction and cooccurrence of colour morphs of the Asiatic golden cat, Bhutan. Cat News 62: 18–20.

Villa, A. G., Salazar, A., & Vargas, F. (2017). Towards automatic wild animal monitoring: Identification of animal species in camera-trap images using very deep convolutional neural networks. Ecological Informatics, 41, 24–32.

Young, S., Rode-Margono, J., & Amin, R. (2018). Software to facilitate and streamline camera trap data management: A review. Ecology and Evolution, 8(19), 9947–9957.

Yousif, H., Yuan, J., Kays, R., He, Z. (2019) Animal Scanner: Software for classifying humans, animals, and empty frames in camera trap images. Ecology and Evolution, 9: 1578–1589.

Weinstein, B. G. (2015). Motion Meerkat: integrating motion video detection and ecological monitoring. Methods in Ecology and Evolution, 6(3), 357–362.

Willi, M., Pitman, R. T., Cardoso, A. W., Locke, C., Swanson, A., Boyer, A., Veldthuis M., & Fortson, L. (2018). Identifying animal species in camera trap images using deep learning and citizen science. Methods in Ecology and Evolution, 10 (1):80–91.

Zhang, J., Wu, X., & Sheng, V. S. (2016). Learning from crowdsourced labeled data: a survey. Artificial Intelligence Review, 46(4), 543–576.

Zhang, X., Yang, W., Tang, X., & Liu, J. (2018). A Fast Learning Method for Accurate and Robust Lane Detection Using Two-Stage Feature Extraction with YOLO v3. Sensors, 18(12), 4308.

## References

Beery, S., Van Horn, G., & Perona, P. (2018). Recognition in terra incognita. In Proceedings of the European Conference on Computer Vision (ECCV) (pp. 456–473).

